# A quantitative imaging framework reveals density-dependent GPCR oligomerization and organization in living cells

**DOI:** 10.64898/2026.05.19.726161

**Authors:** Céline Delaitre, André Dias, Nikolaj Brinkenfeldt, Elisa Pons, Mathew Mungra, Gabriella von Scheel von Rosing, Jacob Hallberg, François Dupuis, Sandra Lecat, Poul Martin Bendix, Morten M. Meldal, Mette M. Rosenkilde, Signe Mathiasen, Karen L. Martinez

## Abstract

GPCR oligomerization has been reported for decades, yet its extent and functional relevance in living cells remain unresolved because existing approaches, often done in bulk, are poorly account for local receptor density, a major determinant of intermolecular interactions. Here, we establish a generic quantitative imaging framework that links spatially resolved FRET measurements describing protein oligomerization to local membrane protein in living cells. Using automated high-throughput analysis of fluorescence images, the method generates large density-resolved datasets that enable direct quantification of receptor oligomerization parameters, including apparent affinity, oligomerization state, and monomer/dimer populations at the submicrometer scale. Applied to class A GPCRs in HEK293 cells, the approach reveals receptor-specific density-dependent equilibria between monomers and dimers over physiologically relevant expression ranges, with no evidence for stable higher-order oligomers under basal conditions. The receptors studied exhibit distinct apparent affinities for dimerization, ranging from predominantly monomeric to dynamic monomer–dimer equilibria, indicating that local membrane density strongly influences receptor organization and that it is receptor dependent. The agreement between our measurements and low-density single-molecule studies further suggests that previously reported higher-order oligomers may partly reflect density-driven receptor proximity effects. By bridging single-molecule and ensemble measurements within a unified quantitative framework, this work reconciles conflicting observations in the GPCR oligomerization literature and provides a broadly applicable strategy for investigating membrane protein organization in living cells.

**Significance:** GPCR oligomerization in living cells is strongly influenced by the local protein density, yet most approaches do not quantitatively account for this parameter. Here, we introduce a quantitative high-throughput imaging framework that directly relates membrane protein local density to local oligomerization state in living cells.

Applied to distinct GPCRs over physiologically relevant density ranges, the method reveals distinct density-dependent monomer–dimer equilibrium and apparent affinities for self-association. These results help reconcile longstanding discrepancies, where distinct oligomerization states have been measured depending on experimental conditions.

More broadly, this work establishes local membrane protein density as a key determinant of membrane protein organization, and provides a quantitative framework applicable to membrane protein complexes in their native cellular context.

## Introduction

G protein–coupled receptors (GPCRs) constitute one of the largest and most diverse families of membrane proteins, playing central roles in cellular communication and physiological processes regulation. They are involved in a wide range of pathological conditions and remain among the most important pharmacological targets to date, accounting for approximately 36% of FDA-approved drugs (1). Through their ability to couple heterotrimeric G proteins and interact with a variety of intracellular partners, GPCRs activate complex and finely tuned signaling networks essential for appropriate cellular responses to external stimuli.

Beyond their canonical signaling behavior, GPCRs have also been identified in various oligomeric assemblies. Evidence for GPCR oligomerization has accumulated over several decades (2-6), revealing that class C GPCRs, like the metabotropic glutamate receptor 2 (mGluR2), exist as constitutive dimers, while many others have been reported to exist as monomers and higher-order oligomers). (4, 7)

GPCR oligomerization has been shown to influence ligand pharmacology (4, 8), both ligand affinity and efficacy (4, 9-14), impose signaling bias (10, 13), and regulate receptor trafficking as well as membrane compartmentalization (15). These observations point to an additional layer of signaling diversity depending on GPCR oligomerization state. Furthermore, assembly of protomers into oligomers provides additional druggable interfaces beyond classical orthosteric sites, an aspect particularly relevant for bitopic and allosteric ligands engineered to achieve pathway, or state-selective modulation (4, 16, 17).

The evidence of the functional role of oligomers, their observations in native tissues, where they have been shown to play important roles in both physiological and pathological processes (4, 18, 19), together with the new perspectives for drug discovery, underscores the need to rigorously examine the mechanisms of oligomer formation and their functional consequences.

Despite this relevance, studies directly addressing the functional roles of GPCR homodimers remain relatively scarce, largely because most cellular systems co-express multiple species of a given GPCR (eg monomers, homodimers, oligomers), making it difficult to attribute specific functional outcomes to defined oligomeric assembly type/species. More fundamentally, key aspects of GPCR homo-oligomerization, including the stoichiometry, interface between protomers, spatial organization, dynamics in living cells, and function, remain unresolved and continue to be actively debated (20-23). For example, several studies have shown conflicting or divergent findings about oligomerization state (both homo- and hetero-) for the same GPCR (24), for example, in the case of D2R (13).

Indeed, reported observations are often highly dependent on the experimental approach employed (25, 26), and the literature frequently reveals limited convergence across methodologies (7, 8). Substantial discrepancies persist between studies, often arising from differences in receptor expression systems, labeling strategies, detection sensitivity, spatial resolution, or data interpretation frameworks, and in some cases leading to conflicting conclusions regarding both the prevalence and functional relevance of GPCR oligomerization (15, 25). It has been the case for the β2-adrenergic receptor (β2AR), for which single-molecule imaging studies have suggested a predominantly monomeric and transiently interacting population (27), whereas ensemble Förster Resonance Energy Transfer (FRET) and Bioluminescence Resonance Energy Transfer (BRET) measurements have supported the existence of stable dimers or higher-order oligomers (28).

Most evidence supporting GPCR oligomerization has been obtained using ensemble RET-based assays, particularly microplate reader-based BRET approaches (29, 30), and FRET-derived methods including HTRF/TR-FRET (31). These approaches have provided important insights into receptor proximity but are subject to limitations that complicate quantitative interpretation. Because these measurements are performed in bulk, the resulting signals reflect an ensemble average, obscuring cell-to-cell variability and heterogeneity in receptor organization. Accurate control of the donor-to-acceptor ratio within the sample remains challenging, despite its strong impact on RET efficiency and, consequently, on the interpretation of oligomeric states (32).

Another important limitation of most current studies is the lack of single cell- and subcellular resolution. Bulk measurements cannot distinguish cells co-expressing both donor- and acceptor-labeled receptors from cells expressing only one (or none) labelled species. Moreover, some ensemble approaches, like BRET, fail to discriminate receptors present at the plasma membrane from intracellular receptor pools. Access to spatial resolution through bioimaging, therefore, constitutes a major advantage for investigating protein–protein interactions, including GPCR homo-oligomerization. By circumventing cellular averaging, image-based approaches provide direct insight into the heterogeneity of receptor organization across individual cells and membrane compartments. In this context, FRET imaging offers unique advantages, and several studies have used it to examine GPCR oligomerization at the single-cell (33, 34) or single proteoliposome level (35, 36).

Moreover, GPCRs are not uniformly distributed across the plasma membrane. They often segregate into microdomains, nanoclusters, or specialized membrane subregions, resulting in local fluctuations in receptor density. These local variations, for example due to lipid domains, are likely to influence the equilibrium between monomeric and oligomeric states, making global averages insufficient to capture true oligomerization behavior (37-40). Single-molecule studies have described the coexistence of monomers, dimers, trimers, and tetramers (20, 41), with the relative abundance of each species strongly influenced by receptor expression levels, and local membrane environment. In addition, transitions between monomeric and dimeric states have been observed by single-molecule FRET microscopy (42), highlighting the dynamic and context-dependent nature of GPCR organization. These results highlight the need of quantitative studies at the submicrometer scale, which will bridge the spatial gap between single-molecule measurements and whole-cell observations.

Quantitative studies at the submicrometer scale appear also relevant because GPCR signaling has been shown to be influenced by GPCR expression levels (43). Variations in receptor abundance may alter receptor proximity and self-association, thereby influencing the composition of signaling complexes and downstream responses (44). However, the relationship between receptor density, oligomerization state, and signaling bias remains poorly defined in intact cells. Thus, our understanding of GPCR oligomerization still lacks a rigorous, quantitative framework operating at the mesoscopic (subcellular) scale that can relate local receptor density to apparent oligomerization state in intact cells.

Here, we present a generic imaging-based approach for the quantitative determination of the apparent oligomerization state of transmembrane proteins within local plasma membrane domains, and apply it to the analysis of GPCR homo-oligomerization. We demonstrate, on several GPCRs, that it enables the link of local receptor densities in living cells to the relative abundance of monomeric and dimeric receptor species. Together, these results provide a unified quantitative framework for interpreting GPCR homo-oligomerization at the mesoscale level, within submicrometer membrane fragments of living cells. Our approach thereby bridges the spatial gap between studies done at the single-molecule and whole-cell levels, and offers mechanistic insight into receptor organization with implications for signaling, biased responses, and pharmacological targeting.

## Results

### 1. Automated line-scan analysis enables high-throughput acquisition of quantitative FRET measurements across heterogeneous subcellular receptor densities

The imaging-based approach exploits the quantitative analyses of confocal fluorescence images of cell populations expressing fluorescently labeled SNAP-tag (ST) GPCRs (ST-GPCRs) with membrane-impermeable donor (D) and acceptor (A) fluorophores (Fig. 1A). This strategy enables receptor labelling with fluorophores, that possess superior optical properties compared to fluorescent proteins, as well as homogenous labeling of the entire cell population with a controlled donor-acceptor ratio (x_D_ = [D]/([D]+[A])) through simple ratiometric mixing of fluorescent substrates. We have optimized labeling conditions to ensure that all receptors at the plasma membrane were labeled, and that the fluorophore stoichiometry matched that of the labeling solution (Fig. S1), which is required to achieve the quantitative labeling suitable for quantitative analysis.

**Figure 1:**
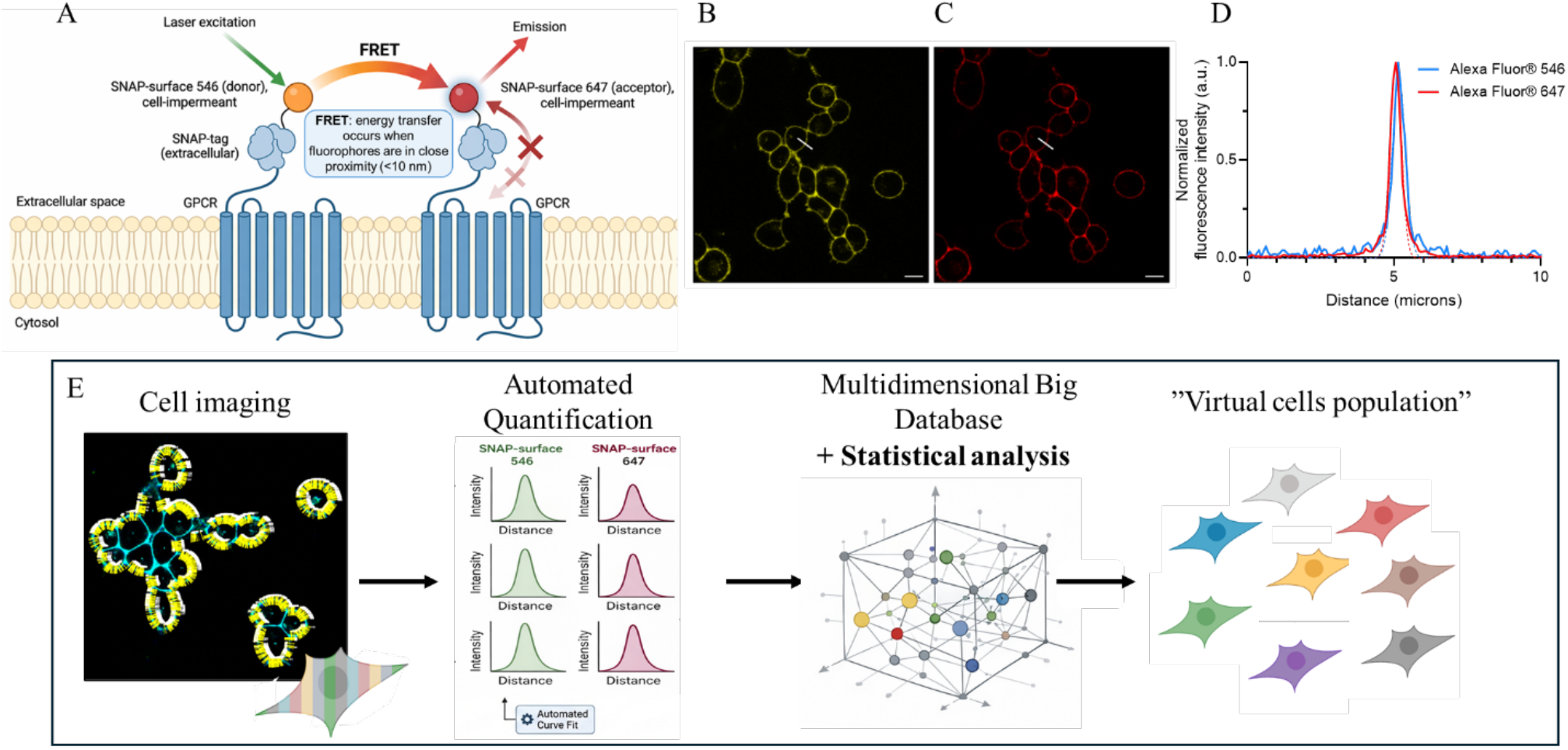
Overview of the FRET-based method for quantifying GPCR oligomerization. (A) Schematic of SNAP-tag labeling strategy used to measure GPCR dimerization by FRET. (B-C) Representative confocal laser scanning microscopy (CLSM) images of HEK293 cells expressing β1AR labeled with donor (B) and acceptor (C) fluorophores. (D) Example of fluorescence intensity profiles extracted from line scans drawn on raw images. (E) Overview of the automated, high-throughput workflow enabling quantitative analysis of membrane fluorescence from multiple line scans, allowing large-scale quantitative analysis of FRET-based measurements. Fluorescence intensities are extracted from numerous local membrane regions within individual cells, which typically exhibit heterogeneous receptor densities (left). These local measurements are pooled and binned according to receptor density, generating “virtual cell populations” with defined and homogeneous density ranges (right), enabling robust statistical analysis across comparable conditions.

We have chosen to use sensitized-acceptor-emission FRET to assess GPCR oligomerization because it directly reports donor–acceptor energy transfer while allowing precise control of donor–acceptor stoichiometry. In contrast to photobleaching- or lifetime-based FRET approaches, sensitized-acceptor-emission FRET limits contributions from nonspecific proximity arising from receptor crowding, membrane microdomain co-localization, or transient encounters, and thereby enables preferential detection of stable and structurally defined receptor–receptor interactions rather than rare or short-lived contacts (45). As a result, it provides a conservative and mechanistically informative readout of oligomerization that minimizes overestimation due to mesoscale membrane organization or expression-dependent clustering. It is thus particularly suitable in the case of class A GPCRs, which are thought to exist predominantly as monomers in dynamic, low-affinity dimerization equilibria. Contributions of nonspecific FRET have been further limited experimentally by performing all our studies with a controlled donor-acceptor ratio (x_D_) comprised between 0.1 and 0.5, for which nonspecific FRET contributions are negligible (45) because donor molecules outnumber acceptor molecules.

Thus, the low FRET efficiencies measured by sensitized-acceptor-emission FRET here, reflect stringent detection criteria rather than a lack of sensitivity.

The automated quantitative analysis of confocal fluorescence images acquired at optimal optical resolution is performed by extracting line scan profiles perpendicular to the cell membrane to quantify local fluorescence signals at multiple locations of the cell membrane (Fig. 1B-D). Depending on the receptor, it corresponds to 11,861-74,066 line scans measured on 42 – 495 distinct cells. Each line scan corresponds to fluorescence intensities for directly excited acceptor (IAA) and donor (IDD), as well as acceptor emission resulting from FRET upon donor excitation (IAD), based on which, local measurements of the apparent FRET efficiency (Eapp,se) can be calculated at multiple locations.

Eapp,se was corrected for nonspecific intermolecular FRET arising from direct acceptor excitation and donor bleed-through using IAD for donor only (x_D_ = 1) and acceptor only (x_D_ = 0) (Fig. S2A-B; Sup. Text). The resulting traces confirm efficient removal of optical crosstalk while preserving density-dependent FRET signals (Fig. S2C).

As shown in the case of the angiotensin II Type 1 receptor (ST-AT1R) (Fig. S3A-B), we observed, for all the receptors studied, a broad distribution of IAA, which reveals inherent heterogenous receptor distributions at the plasma membrane within cells (Fig. S3B) and between cells (Fig. S3A). Total receptor intensity (RI) was estimated from IAA by accounting for x_D_, thereby enabling accurate quantification of the total receptor population at the plasma membrane, which can later on be converted into molar densities using conversion factors determined by the Calmet method for the imaging settings used (46).

A representative selection of these measurements illustrates the relationship between apparent FRET efficiency and receptor density across a broad range of expression levels reached according to the transient transfection conditions used (Fig. S3C). For each of the three independent experiments (minimum), receptor intensities were divided into bins of comparable densities (with more than 100 measurements per density range) (Fig. S3D), thereby providing strong statistical power for subsequent analysis. Each of the bin, is used to generate “virtual cells”, defined as computationally reconstructed populations with homogeneous receptor density ranges (Fig. 1E).

The statistical robustness of the binning approach was confirmed by verifying that receptor intensity measurements remain consistent across different levels of analysis (Fig. S3E). The median receptor intensity is comparable when calculated across the entire dataset, individual frames, cells, or local line scans, indicating that each bin represents a statistically relevant subset of the data. This demonstrates that the line-scan approach does not introduce bias in the quantification, while the very low SEM values observed (Fig. S3F) further indicate that this strategy enables accurate intensity measurements. In the case of AT1R, these virtual cells exhibit receptor density variations typically below 15% Fig. S3E), producing measurements that are substantially more homogeneous than the stochastic fluctuations expected within a single cell (47).

### 2. AT1R exhibits density-dependent homo-dimerization at the plasma membrane

In the case of ST-AT1R, Eapp,se, measured for labeled samples (x_D_ = 0.5), showed a clear density-dependent increase, whereas a negligible Eapp,se, close to baseline was detected across the expression range allowed in the case of Δ2Δ for the best transient transfection conditions found, an established monomeric control, composed of the transmembrane fragment of mGluR2 lacking its amino- and carboxy-terminal domains (20, 48) (Fig. 2A).

**Figure 2:**
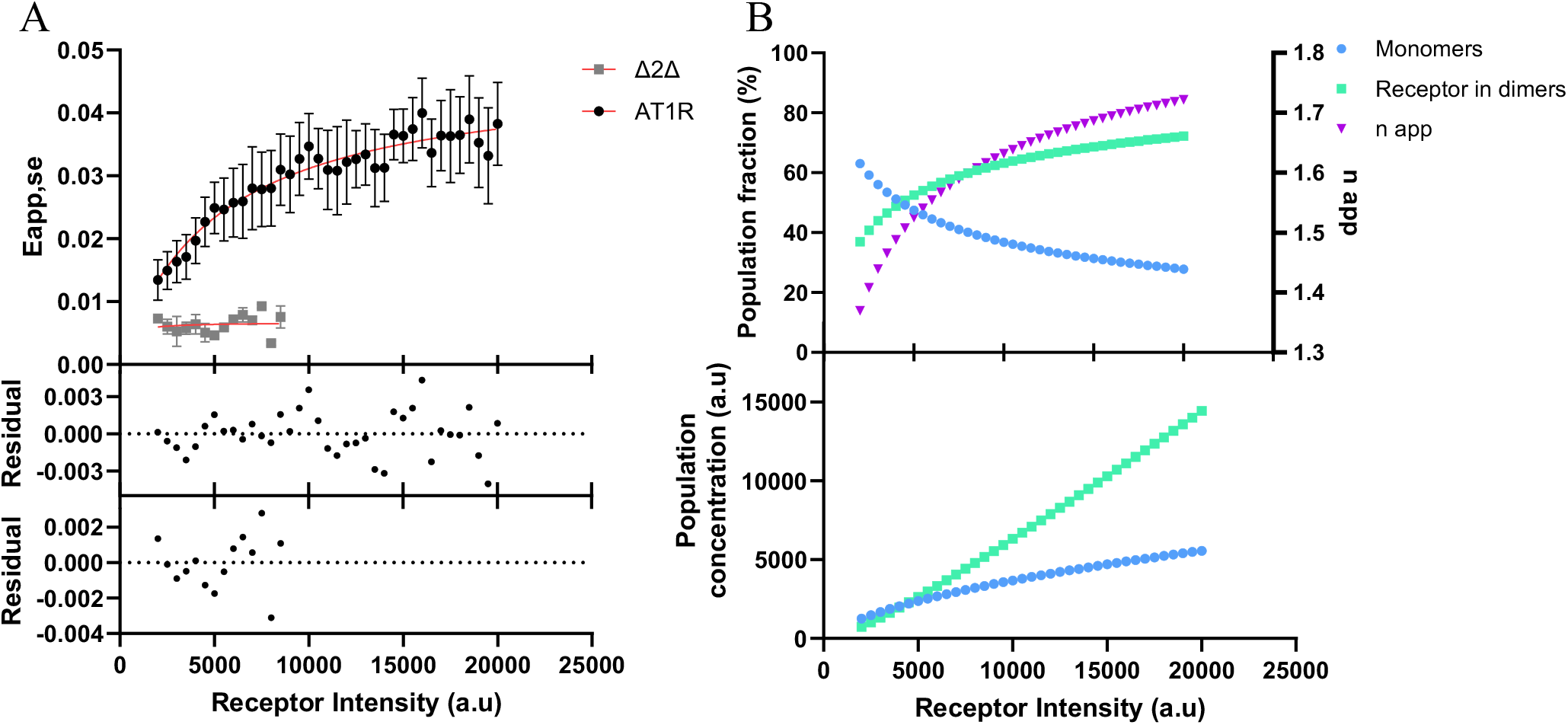
The oligomerization state of AT1R depends on receptor density in HEK293 cells. (A) Apparent FRET efficiency (Eapp,se) plotted as a function of receptor surface density (arbitrary units) in HEK293 cells transiently expressing SNAP-AT1R or the SNAP-Δ2Δ control, measured at a donor mole fraction (x_D_) of 0.5. Each point represents the mean ± SEM obtained from n = 4 independent experiments for AT1R and n = 2 for Δ2Δ. Data were fitted using the one-site specific binding model (GraphPad Prism), yielding an apparent dissociation constant (K_d_) of 5049 a.u (31.1 proteins/µm^2^) and an X^2^ of 0.01405. Residuals of the fit are shown below the plot. (B) Using the K_d_ value derived from the fit in A, the fractions of monomeric and dimeric receptors were estimated and plotted as percentages as a function of receptor surface density (left y-axis), while the corresponding apparent oligomerization number (n_app_) is shown on the right y-axis. The corresponding concentrations of monomeric and dimeric receptor species as a function of receptor surface density are shown directly below.

We used the residual fluctuations in Eapp,se of this pure monomeric protein Δ2Δ to estimate the intrinsic experimental noise in our quantitative studies. The dispersion of Eapp,se values across the explored density range yielded a root mean square error (RMSE) of 0.0019, defining the effective noise baseline in our measurements, below which variations in apparent FRET efficiency cannot be distinguished from background fluctuations. Assuming normally distributed errors, the practical detection limit corresponds to approximately three times the RMSE (Eapp,se ≈ 0.0056), such that only density-dependent increases exceeding this threshold can be reliably attributed to receptor–receptor interactions. This provides an objective criterion for interpreting low FRET signals, and highlights that the absence of detectable density-dependent FRET does not necessarily imply the absence of receptor interactions, but rather that such interactions fall below the sensitivity limit within the explored density range.

We evaluated several models to describe the oligomerization behavior of ST-AT1R (see Material & methods). The data were best fit by a simple model—previously applied in single-molecule studies (27), in which monomeric receptors progressively assemble into dimers as receptor density increases. This system can be quantitatively described using a one-site specific binding model (Strategy 1), which yielded an apparent homodimerization constant K_d_ of approximately 5.0 x 10^3^ a.u. (i.e. 30.9 proteins/µm^2^) (46) that can be used to determine the fractions of monomeric and dimeric receptors, their respective concentrations, and the apparent oligomerization number (n_app_) (Fig. 2B).

As receptor density increases, the fraction of monomeric receptors progressively decreases while the fraction of receptors in dimers increases until reaching an equilibrium between monomers and dimers (Eapp_max)_. Consequently, n_app_ increases until reaching ∼1.7 for receptor densities above 15 000 a.u. (i.e. 83.7 proteins/µm^2^), while the absolute concentrations of receptors in both monomeric and dimeric forms continue to increase with receptor expression.

For AT1R, we show that the large datasets obtained upon image analysis enable the extraction of a complete set of quantitative parameters describing receptor oligomerization, including the apparent K_d_, the maximum FRET efficiency (Eapp_max)_, as well as parameters varying for each receptor density, such as n_app_, densities, and relative fractions of monomers and dimers.

### 3. Parameter uncertainty analysis confirms robustness of AT1R oligomerization estimates

We next examined the robustness of our analyses and the influence of experimental variability on the estimation of the apparent K_d_ and the resulting receptor populations. Independent fits of four AT1R datasets yielded comparable yet distinct K_d_ values (Fig. S4A), resulting in modest differences in the predicted monomer and dimer fractions (Fig. S4B). To further quantify errors for K_d_, monomer, and dimer fractions, Monte Carlo simulations were performed by generating 1000 synthetic datasets incorporating the experimentally determined noise level. Refitting these datasets produced a distribution of K_d_ values centered on the experimentally determined mean, with a 99% confidence interval of approximately 3.9–6.6 x 10^3^ a.u. (Fig. S4C). Propagation of this uncertainty to receptor populations showed that the inferred monomer and dimer fractions remain tightly constrained across the explored range of receptor densities (Fig. S4D), with variations within ∼5%, indicating that the conclusions regarding AT1R oligomerization are robust to experimental noise and parameter uncertainty.

### 4. Not all class A GPCRs display similar density-dependent oligomerization at the plasma membrane

To determine whether density-dependent oligomerization is a general feature among GPCRs, we applied the same analysis to several class A receptors (Fig. 3A-B). Among the receptors tested, C-X-C motif chemokine receptor 4 (ST-CXCR4), beta-1-adrenergic receptor (ST-β1AR) and beta-2 adrenergic receptor (ST-β2AR) displayed distinct density-dependent increases in Eapp,se above the experimental noise level (Fig. 3A-B), allowing the use of the one-site specific binding model as done in the case of ST-AT1R to determine apparent homodimerization constants (K_d_) and the corresponding monomeric and dimeric fractions and concentrations, and n_app_ (Fig. S5, Table 1).

**Table 1:** Summary of parameters derived from density-dependent FRET analysis of GPCR oligomerization.

| Receptor | Class | $K_d$ (a.u.)<br>(i.e. $p/\mu m^2$) | $E_{app_{max}}$ | R 50 (a.u.)<br>(i.e. $p/\mu m^2$) | Req (a.u.)<br>(i.e. $p/\mu m^2$) | $n_{app_{max}}$ |
| --- | --- | --- | --- | --- | --- | --- |
| AT1R | A | $5049 \pm 1272$<br>($31.13 \pm 11.18$) | $0.047 \pm 0.004$ | $\sim 5000$<br>($\sim 30.88$) | $\sim 10000$<br>($\sim 57.29$) | $\sim 1.7$<br>($\sim 1.85^*$) |
| $\beta 1AR$ | A | $5686 \pm 2151$<br>($34.50 \pm 15.82$) | $0.032 \pm 0.004$ | $\sim 5500$<br>($\sim 33.52$) | $\sim 9000$<br>($\sim 52.01$) | $\sim 1.69$<br>($\sim 1.8^*$) |
| $\beta 2AR$ | A | $6979 \pm 2982$<br>($41.33 \pm 20.21$) | $0.017 \pm 0.003$ | $\sim 7000$<br>($\sim 41.44$) | $\sim 12000$<br>($\sim 67.85$) | $\sim 1.66$<br>($\sim 1.76^*$) |
| CXCR4 | A | $1263 \pm 373$<br>($11.13 \pm 6.43$) | $0.048 \pm 0.002$ | $< 2000$<br>($< 15.03$) | $\sim 5000$<br>($\sim 30.88$) | $\sim 1.84$<br>($\sim 1.9^*$) |
| CB1R | A | $<<$ | $0.013 \pm 0.002$ | ND | ND | $\sim 1.21^*$ |
| 5-HT2AR | A | $<<$ | - | ND | ND | $\sim 1.13^*$ |
| $\Delta 2\Delta$ | C | $<<$ | $0.007 \pm 0.001$ | ND | ND | $\sim 1.07^*$ |
| mGluR2 | C | $>>$ | $0.083 \pm 0.002$ | ND | ND | $\sim 1.93^*$ |

a.u.: arbitrary unit; -: Nonconvergent $K_d$ values indicate the absence of detectable density-dependent population changes within the explored expression range, consistent with predominantly monomeric or dimeric receptor populations; R 50: Receptor Intensity when 50% Monomers / 50% Dimers; Req: RI when equilibrium is reached; ND: not determined \*: determined with Strategy 2

**Figure 3:**
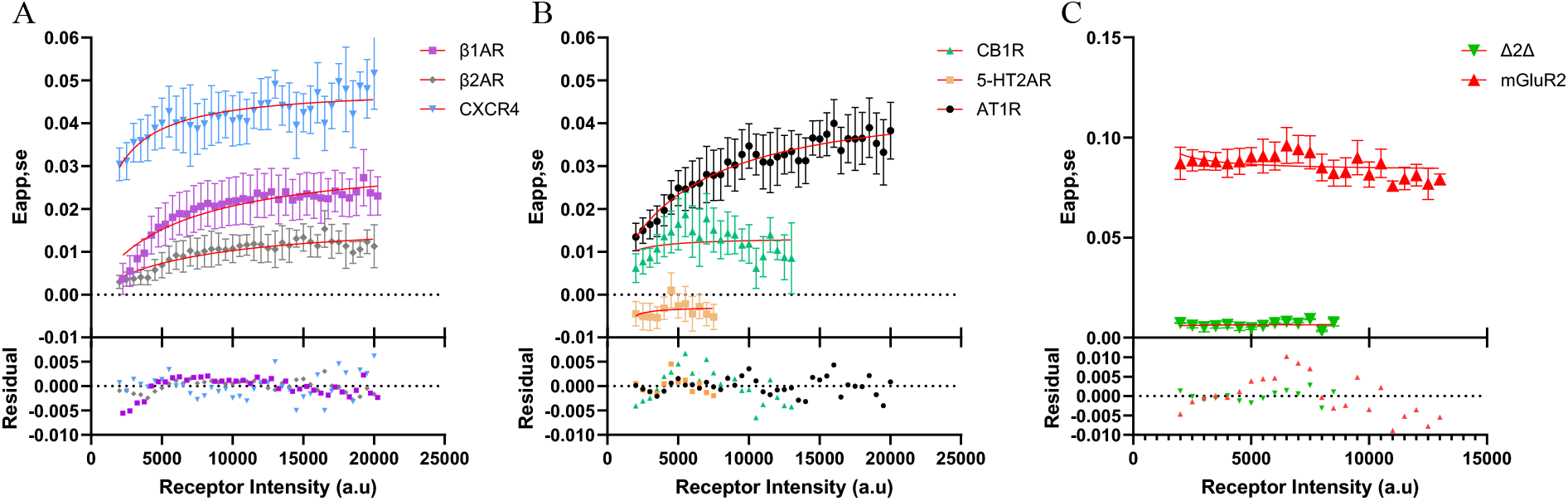
Comparative analysis of density-dependent oligomerization across class A GPCRs and controls. (A-C) Eapp,se plotted as a function of receptor surface density (a.u.) in HEK293 cells transiently expressing SNAP-tagged class A GPCRs, including β1AR, β2AR, CXCR4, AT1R, CB1R, and 5-HT2AR (A-B) or mGluR2 and Δ2Δ (C). Symbols represent mean ± SEM of experimental data, and solid lines indicate nonlinear regression using the One-site specific binding model described in Fig. 2 (GraphPad Prism). Residuals of the fit are shown below the plots. All receptors were analyzed using a minimum of n = 3 independent experiments except Δ2Δ (n = 2).

In contrast, cannabinoid receptor 1 (ST-CB1R) and 5-hydroxytryptamine receptor 2A (ST-5-HT2AR) displayed very low, constant, Eapp,se across the explored receptor density range, suggesting low, or no, constant oligomerization rate (Fig. 3B). Notably ST-5-HT2AR values are lower than those obtained with the truncated Δ2Δ construct, which served as a monomeric control and defines the biological noise level of the method determined before (RMSE Eapp,se ≈ 0.0056) (Fig. 3C). This indicates that 5-HT2AR falls within the noise range associated with non-specific proximity, preventing reliable quantification of oligomerization using Strategy 1 (density-dependent Strategy).

A density-independent, but high Eapp,se value was observed for ST-mGluR2, which indicates a strong oligomerization rate between GPCRs, that we cannot determine with the range of expression levels available in our data, which would be consistent with its constitutive oligomerization (49)(Fig. 3C).

In such cases, a complementary analysis based on the dependence of Eapp,se on x_D_ is employed (Strategy 2).

### 5. Quantification of receptor oligomerization using donor-acceptor ratio analysis

The accurate control of x_D_ allowed us to implement a second Strategy based on the dependence of apparent FRET efficiency on x_D_ (34, 45), where n_app_ is directly determined using the theoretical framework established by Meyer *et al*. (34). To implement this approach, datasets at multiple x_D_’s were required. To avoid proximity-induced FRET (Eprox), x_D_ values were restricted to ≤ 0.5, ensuring that Eapp,se reflects specific receptor interactions rather than nonspecific proximity effects (45).

This Strategy was first validated on AT1R. Fitting of traces Eapp,se = f(x_D_) with equation 5 (Fig. 4A; see methods), allowed the determination of specific n_app_ for each range of expression analyzed. n_app_ revealed a progressive shift of the receptor population from monomers toward dimers as receptor density increases (Fig. 4B). Importantly, the values of n_app_ are in good agreement with those obtained with the first Strategy, corresponding to the density-dependent Strategy (Fig. 4C), thereby confirming that our two analytical strategies both provide a quantitative determination of n_app_ at the sub-micrometer scale.

**Figure 4:**
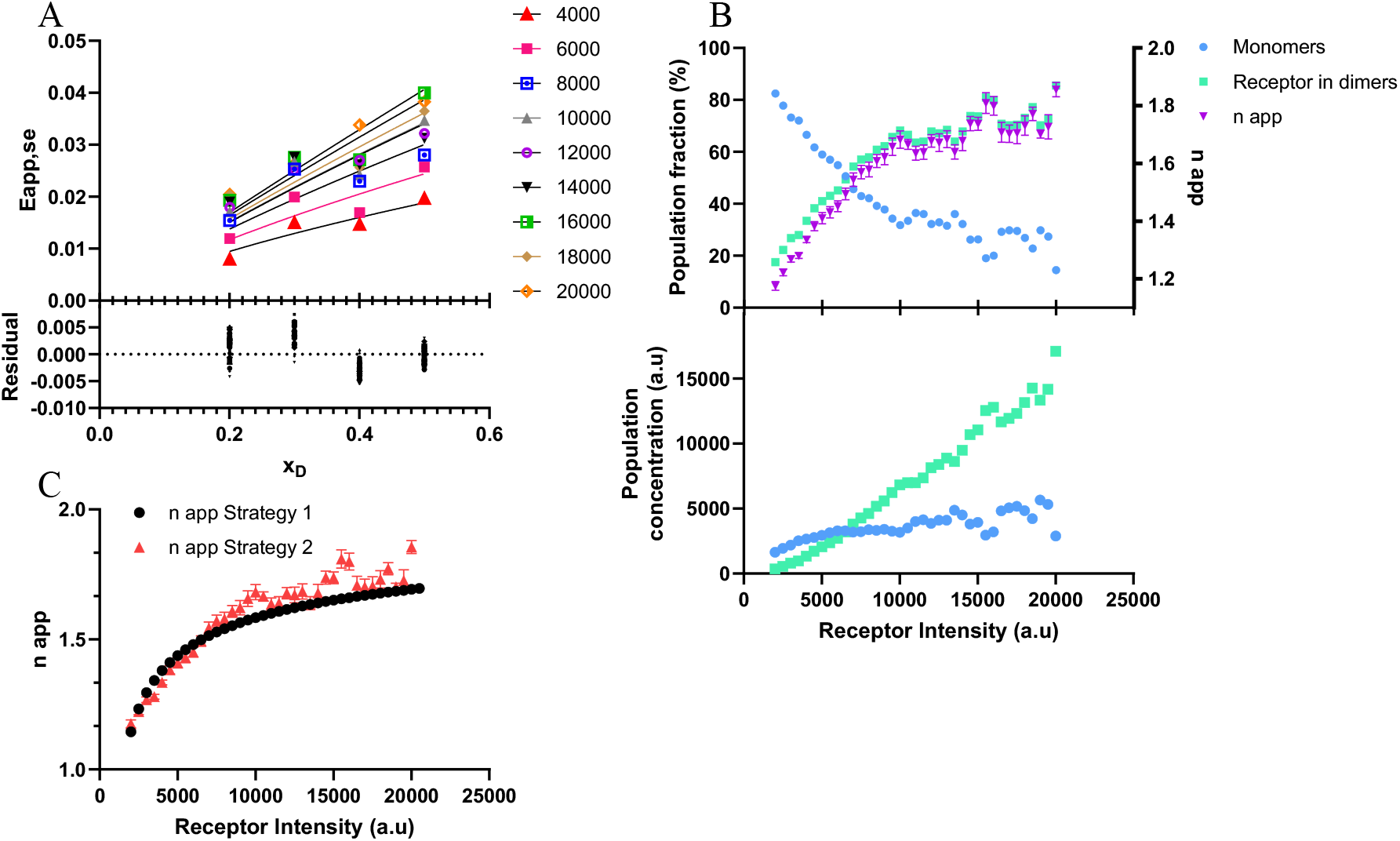
Estimation of AT1 receptor oligomerization at surface density using donor fraction dependence. (A) Apparent FRET efficiency (Eapp,se) measured as a function of donor mole fraction (x_D_) at several receptor surface densities in HEK293 cells transiently expressing ST-AT1R. Data were fitted using a second oligomerization model (Eq. 5; GraphPad Prism). Residuals of the fits are shown below. (B) Apparent oligomerization number (n_app_) estimated from the donor fraction–dependent analysis (right y-axis), together with the corresponding fractions of monomeric and dimeric receptors plotted as percentages as a function of receptor surface density (left y-axis). (C) Comparison of n_app_ values obtained from the Strategy 1 (surface density–dependent analysis) and from the Strategy 2 (donor fraction–dependent analysis).

The determined n_app_ values for CXCR4, β1AR, and β2AR with Strategy 2 (Table 1) were consistent with those obtained with Strategy 1, confirming that these receptors exist in dynamic monomer–dimer equilibria at the plasma membrane (Fig. 5).

**Figure 5:**
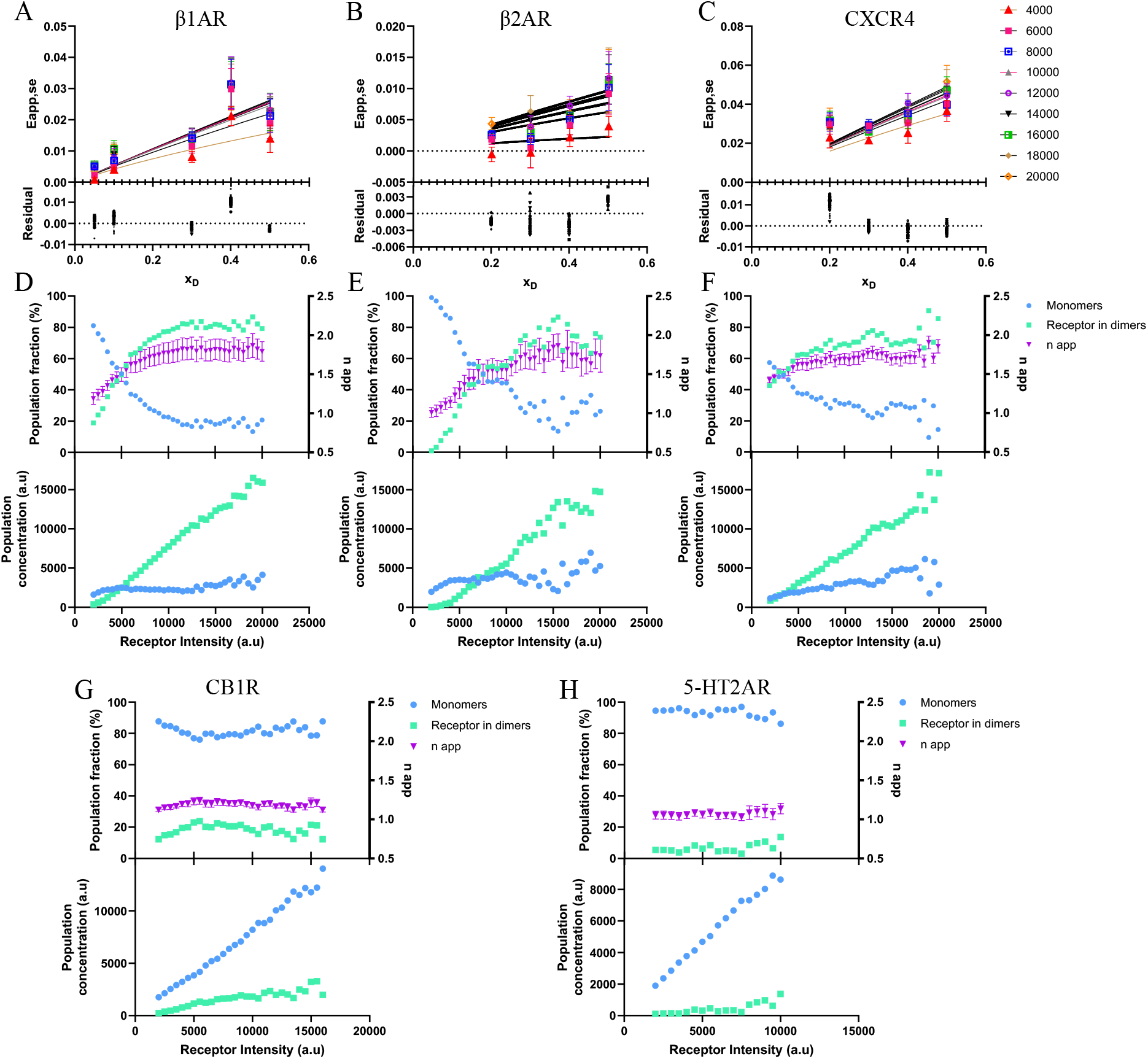
Estimation of class A receptor oligomerization at surface density using donor fraction dependence. (A-C) Apparent FRET efficiency (Eapp,se) measured as a function of donor mole fraction (x_D_) at several receptor surface densities in HEK293 cells transiently expressing ST-β1AR (A), ST-β2AR (B) or ST-CXCR4 (C). Data were fitted using a second oligomerization model (Eq. 5; GraphPad Prism). Residuals of the fits are shown below. (D-H) Apparent oligomerization number (n_app_) estimated from the donor fraction–dependent analysis (right y-axis), together with the corresponding fractions of monomeric and dimeric receptors plotted as percentages as a function of receptor surface density (left y-axis). Calculations were performed for β1AR (D), β2AR (E), and CXCR4 (F) using the n_app_ values obtained from the second oligomerization model (Eq. 5; GraphPad Prism). Calculations were performed for CB1R (G) and 5-HT2AR (H) using the Etrue from AT1R to obtain the n_app_ values used for the second oligomerization model (Eq. 5; GraphPad Prism). The reported n_app_ values were derived from at least three independent experiments.

The constant low Eapp,se observed for Δ2Δ, CB1R, and 5-HT2AR corresponds to constant n_app_ values close to 1 (Fig. 5G-H, fig. 6), indicating predominantly monomeric organization under the experimental conditions tested. As expected, analysis of mGluR2 yielded n_app_ values close to 2 at any expression levels tested, consistent with its well-established constitutive dimeric organization (49).

**Figure 6:**
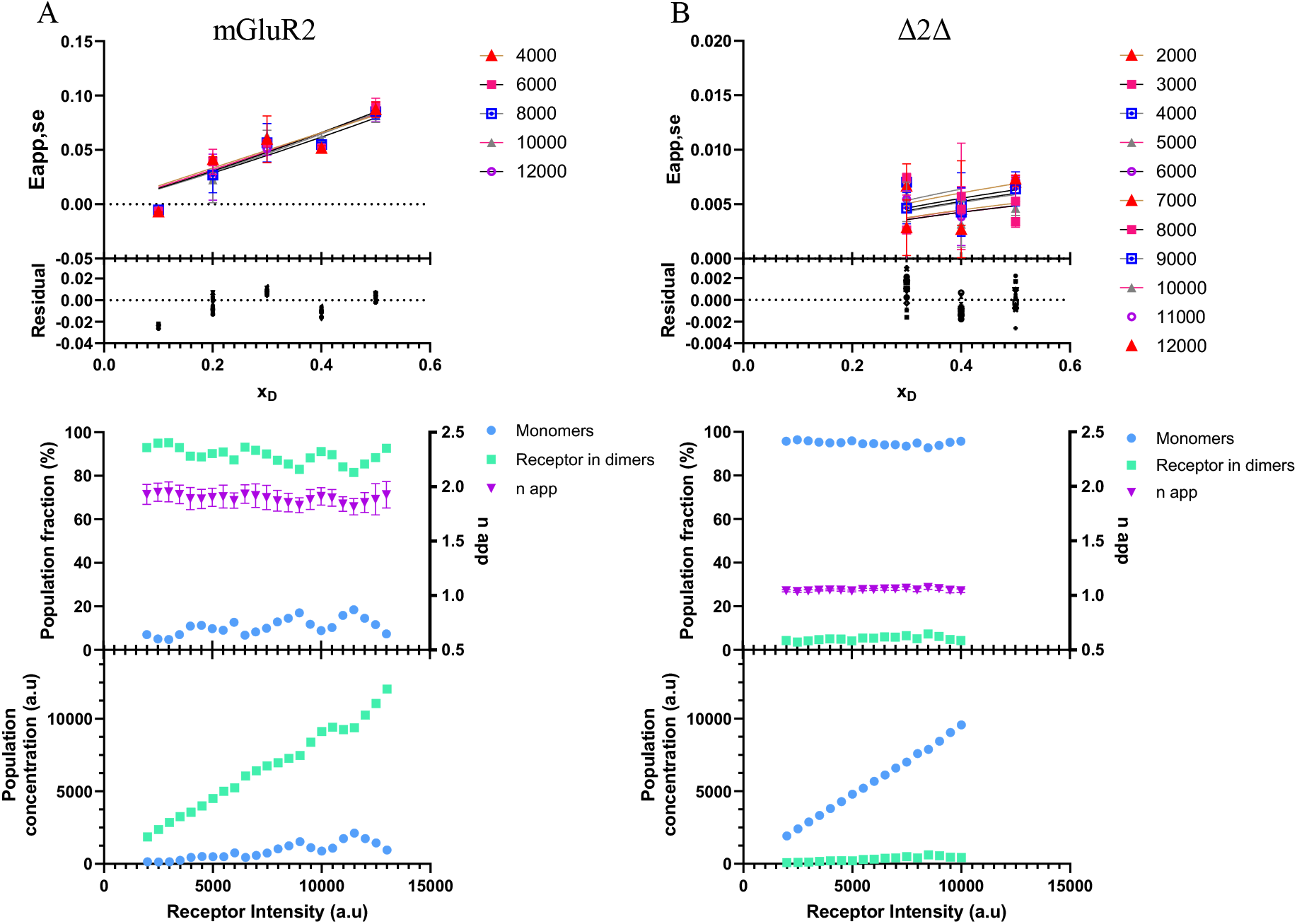
Estimation of class C receptor oligomerization at surface density using donor fraction dependence. (A-B) Apparent FRET efficiency (Eapp,se) measured as a function of donor mole fraction (x_D_) at several receptor surface densities in HEK293 cells transiently expressing ST-mGluR2 (A) or ST-Δ2Δ (B). Data were fitted using a second oligomerization model (Eq. 5; GraphPad Prism). Residuals of the fits are shown below. Data corresponds to n = 4 and n = 2 independent experiments. (C-D) Apparent oligomerization number (n_app_) estimated from the donor fraction–dependent analysis (right y-axis), together with the corresponding fractions of monomeric and dimeric receptors plotted as percentages as a function of receptor surface density (left y-axis). Calculations were performed for mGluR2 using the n_app_ values obtained from the second oligomerization model (Eq. 5; GraphPad Prism). Calculations were performed for Δ2Δ (D) using the Etrue from mGluR2 to obtain the n_app_ values used for the second oligomerization model (Eq. 5; GraphPad Prism).

From the combined use of density-dependent and x_D_–based analyses, it is possible to extract key parameters including the apparent homodimerization K_d_, maximal FRET efficiency (Emax), n_app_, monomeric and dimeric fractions and concentrations, as well as intrinsic FRET efficiency (Etrue). These parameters provide complementary insights into receptor organization, ranging from interaction strength to structural and spatial characteristics of receptor assemblies. A summary of all quantifiable parameters obtained for each receptor is provided in Table 1.

Those two strategies are complementary but with a different degree of uncertainty. While the fit of the curve Eapp,se = f(R) can be done on a large number of data points in the Strategy 1, the analysis in this second Strategy is based on several fits of curves Eapp,se = f(xD), which contain considerably fewer points (between 4 – 6), requires the estimation of Etrue, the FRET efficiency of a fully interacting D-A pair. This makes the data analysis of this second Strategy more sensitive to experimental noise, compared to Strategy 1. Moreover, it also requires more data (the same study for multiple x_D_s) compared to Strategy 1 which only requires only the study at x_D_ = 0.5.

The quantitative evaluation of the uncertainty analysis for n_app_, fractions, and densities of monomers and dimers (Fig. S6) indicated high accuracy for both strategies, with relative uncertainties of approximately 1–2% and 5–7%, respectively, for Strategies 1 and 2. Although the variability associated with Strategy 2 was approximately three-to four-fold larger than that of Strategy 1, it provides a valuable complementary quantitative approach to Strategy 1, thereby enabling the estimation of all values of n_app_, fractions, and densities of monomers and dimers depending on the local receptor density.

## Discussion

We present here a quantitative FRET-based approach to elucidate membrane protein homo-oligomerization at the submicrometer scale. It exploits automated high-throughput analysis of fluorescence images to generate large datasets. The robustness of our approach is due to its scale and high statistical power enabled by the large datasets generated, which allows the partitioning of the cell sample into multiple (at least 30) homogeneous virtual cells and thereby enables highly accurate quantification of oligomerization as a function of receptor densities. This is further strengthened by the establishment of an objective criterion for interpreting low FRET signals, based on the direct estimation of experimental noise and detection limits using a strictly monomeric Δ2Δ construct to define a practical detection threshold.

These large datasets can be analyzed statistically using two complementary strategies. The density-dependent saturation analysis (Strategy 1), used when FRET varies depending on receptor density, provides the most accurate estimates of oligomerization parameters, whereas the x_D_–dependent analysis (Strategy 2) remains valuable and informative in the case of constant FRET signals for the range of receptor densities studied. These constant FRET signals could reflect no or very low affinity between protomers when Eapp,se is low, or high affinity between protomers when Eapp,se is high. Strategy 2 thereby broadens the range of receptors that can be quantitatively analyzed and reduces the risk of overlooking biologically relevant, but low-amplitude interactions. Altogether, the two signal analysis strategies provide, via quantitative oligomerization parameters determined at the virtual cell level, complementary insights into receptor organization at the submicrometer scale, ranging from interaction strength to structural and spatial characteristics of receptor assemblies.

Bulk RET-based studies first suggested that GPCR oligomerization depends on receptor expression level, as variations in DNA transfection were correlated with changes in the averaged apparent oligomerization parameter n_app_ (7, 28). However, these approaches provided only population-averaged measurements corresponding to a single mean expression level for the entire cell sample (7, 50). In contrast, single-molecule methods and subcellular FRET approaches enabled the direct observation of receptor interactions with high spatial precision, and demonstrated the existence of monomer–dimer equilibria at low receptor densities (20, 27, 34, 41, 51-54). Nevertheless, these methods are generally restricted to very low expression levels and do not establish a quantitative framework linking receptor density to oligomerization state across a broad range of membrane densities with submicrometer resolution.

Our approach addresses this gap by establishing a direct quantitative relationship between local receptor density and oligomerization parameters, including Kd, napp, and density-dependent monomer and dimer fractions, over a broad range of expression levels. In this context, it bridges low-density single-molecule regimes and higher-density bulk measurements, while maintaining subcellular spatial resolution. Importantly, the receptor densities explored here span regimes ranging from predominantly monomeric populations to monomer–dimer coexistence and approaching equilibrium saturation. The monomer–dimer equilibria observed across this range are consistent with results obtained by single-molecule FRET studies at low expression levels (20, 27, 51), as well as with observations made in reconstituted membrane systems at physiological densities (51). Moreover, the expression levels obtained under standard transient transfection conditions overlap with those commonly encountered in heterologous systems used in pharmacological assays, suggesting that the equilibria described here, in the absence of ligand, are directly relevant to the interpretation of many commonly used GPCR experimental systems (55-58).

Within this quantitative density-resolved framework, the class A GPCRs analyzed here predominantly exhibit monomer–dimer equilibria across the explored range of receptor densities. AT1R, β1AR, β2AR, and CXCR4 display dynamic density-dependent equilibria between monomeric and dimeric states, consistent with intermediate apparent affinities and with previous single-molecule studies reporting transient dimer formation for these receptors (37, 59, 60). In contrast, CB1R and 5-HT2AR remain predominantly monomeric under similar conditions, with dimer populations below the detection threshold, in agreement with reports describing weak or undetectable dimerization depending on the experimental context (61-64). These observations indicate that receptor-specific apparent affinities strongly influence the extent to which oligomerization varies with local membrane density.

Notably, none of the receptors analyzed here displayed apparent oligomerization states significantly above dimers, despite the broad range of receptor densities explored. Reports of napp values greater than 2 have mainly originated from bulk RET approaches or single-molecule counting techniques (8, 17, 41), which may be influenced by receptor proximity effects and do not necessarily distinguish specifically assembled oligomers from closely apposed receptors at high local density (65). In contrast, single-molecule FRET studies, which are better suited to resolving direct receptor interactions, have predominantly reported monomer–dimer equilibria, although typically at much lower receptor densities (generally around 0.15–0.30 receptors/µm^2^) (20, 41). The agreement between these studies and our results obtained over a substantially broader density range suggests that previously reported higher-order oligomers may, at least in part, reflect density-driven receptor proximity rather than stable higher-order oligomers. Together with structural and biophysical studies indicating that class A GPCR dimers are generally transient and stabilized by weak interfaces (20, 40, 51, 60, 66, 67). Our results support a model in which basal GPCR organization under the conditions tested is dominated primarily by monomer–dimer equilibria.

More broadly, these findings provide a framework for reconciling discrepancies in the GPCR oligomerization literature by demonstrating that oligomerization behavior is strongly dependent on local receptor density. Variations in receptor expression levels across tissues, experimental systems, or pathological conditions are therefore likely to influence receptor organization and contribute to apparent differences in reported oligomeric states, particularly for receptors with intermediate apparent affinities. Because oligomerization equilibria depend on local membrane density, the density at which monomeric and dimeric populations equilibrate may represent a useful parameter for predicting receptor behavior in different cellular contexts and for interpreting variability in pharmacological and functional assays. In addition, as suggested by single-molecule studies (27), these equilibria are likely to shift upon ligand binding in a receptor- and ligand-dependent manner, with potential consequences for GPCR signaling and pharmacology.

An important feature of our approach is the ability to convert fluorescence intensities into absolute receptor surface densities (∼3 to 110 receptors/µm^2^ (44)), thereby enabling direct comparison across experiments and linking imaging-based oligomerization measurements to receptor abundance estimates obtained by biochemical or radioligand approaches (4). Reported physiological GPCR expression levels for well-characterized systems such as β-adrenergic receptors typically correspond to approximately 10^4^–10^5^ receptors per cell, which is expected to translate into surface densities ranging from a few to several tens of receptors per µm^2^ depending on cell size and membrane organization (4, 41). Other receptors, including CXCR4, CB1R, and 5-HT2AR, exhibit substantial variability in expression depending on cell type, membrane environment, and physiological state (61, 63, 68), and are often characterized by heterogeneous or locally enriched membrane distributions. Although physiological receptor densities vary considerably across biological contexts, expressing oligomerization as a function of absolute receptor density provides a framework for evaluating whether the observed monomer–dimer equilibria occur within biologically relevant regimes.

More broadly, access to receptor densities and oligomerization-related parameters (Kd, R50, Req) in absolute rather than arbitrary units enables a more quantitative characterization of receptor organization across physiologically relevant expression ranges. This quantitative framework facilitates objective comparisons between receptors, experimental conditions, and laboratories, independently of the precise distribution of expression levels generated by transient transfection. By linking oligomerization behavior directly to receptor abundance, the approach also provides a common basis for interpreting and comparing GPCR organization across imaging, biochemical, and pharmacological studies.

Beyond oligomerization state, our approach also provides insight into receptor proximity at the structural level. E_true_ provides useful information on receptor proximity and can, in favorable cases, be related to D-A distance. However, structural interpretation of these values should be made with caution. The receptors analyzed here differ markedly in N-terminal length, which is expected to influence fluorophore positioning relative to the transmembrane core and therefore the measured FRET efficiency. This is particularly relevant for receptors such as CB1R and 5-HT2AR, for which the absence of a detectable increase in Eapp,se under the conditions tested limits the extraction of Etrue using Strategy 1. In addition, GPCR dimers are not necessarily restricted to a single rigid geometry, but may adopt multiple interfaces or conformational arrangements (69, 70), making Etrue better interpreted as an effective parameter reflecting an ensemble of receptor configurations rather than a unique structural distance.

Finally, although the present study focuses on homodimerization, many GPCRs are known to form hetero-oligomers with important roles in signaling specificity and pharmacology (3, 4, 13, 57, 71, 72). Extending this quantitative method to heteromeric systems represents an important next step. This approach is not limited to GPCRs but extends to other classes of membrane proteins, offering a framework for linking local protein density to oligomerization state in living cells, to investigate the organization of membrane signaling complexes (73, 74).

## Material and Methods

### Plasmids

Receptors were expressed in HEK293 cells using the pcDNA3.1 mammalian expression vector. All constructs contained the first-generation SNAP-tag. The ST-AT1R, ST-β1AR, and ST-β2AR expression constructs were obtained from Cisbio Bioassays. A ST-CB1R expression construct was generated from the ST-β1AR construct and a CB1R-containing construct acquired from cdna.org (catalog no. #CNR0100001) by restriction cloning. A ST-5-HT2AR expression construct was generated through similar restriction cloning from the ST-β1AR construct and a 5-HT2AR-containing construct provided by Alexander Hauser. Finally, plasmids encoding SNAP-tagged CXCR4 and ST-mGluR2 & ST-Δ2Δ were generously provided by Martin Gustavsson and Jonathan Javitch, respectively.

### Cell lines

Immortalized human embryonic kidney cells HEK293 (ATCC catalog no. CRL-1573, RRID: CVCL_0045) were used and genetically modified with different plasmids according to the different experiments. Flp-In™/T-Rex 293 cells from Invitrogen stably expressing TetOn inducible SNAP-β1AR were established by co-transfection with an excess of pOG44 plasmid encoding Flp recombinase and pcDNA™ 5/FRT/TO/SNAP-β1AR expression vectors.

### Cell culture

HEK293 cells were cultured under standard conditions (37°C, 5% CO_2_) in DMEM (Gibco) supplemented with 10% FBS (Gibco) and 1% antibiotics (100 μg/mL penicillin/streptomycin; ThermoFisher Scientific). To maintain stable plasmid expression in iSNAP-β1AR cells, the medium was supplemented with 15 µg/mL blasticidin S HCl and 100 µg/mL hygromycin B (ThermoFisher Scientific) instead of the 1% antibiotic mixture.

### Expression of SNAP-GPCRs in cells

Approximately 30,000 cells/well were seeded into 8 well-chambered coverslips, precoated with 50µg/mL poly-L-lysine for 15 minutes at room temperature, and incubated overnight at 37°C, 5 % CO_2_. Cells were transiently transfected with ST-GPCR plasmids using Lipofectamine 2000 (ThermoFisher Scientific) according to the manufacturer’s instructions. Briefly, transfection complexes were prepared in Opti-MEM using 1:1 DNA:Lipofectamine 2000 ratio (0.4µg plasmid DNA, 0.4µL lipofectamine in 20µL Opti-MEM per well) and added to wells containing 200µL culture medium. Cells were incubated for 18-24h at 37°C, in 5 % CO_2_. For the iSNAP-β1AR cell line, instead of transfection, receptor expression was induced with tetracycline (1µg/mL; ThermoFisher Scientific). For HEK293 cells transfected with ST-mGluR2, 6 hours after transfection, the medium is replaced with glutamine-free DMEM medium (Gibco).

### Covalent labeling of GPCRs coupled with-tag® by SNAP-Surface Alexa Fluor® 546 and SNAP-Surface Alexa Fluor® 647

Cells expressing ST-GPCRs were labeled with 2.5 µM fluorophore mixture (1.25µM SNAP-Surface Alexa Fluor® 647 and 1.25µM SNAP-Surface Alexa Fluor® 546 for x_D_ = 0.5) in imaging buffer (145 mM NaCl, 5 mM KCl, 1 mM MgCl_2_, 1 mM CaCl_2_, 10 mM HEPES, 10 mM Glucose, pH 7.2) by covalent SNAP-tag coupling for 30 minutes at 37°C. Fluorophore stock concentrations and donor-acceptor mixing ratios were verified by absorbance spectroscopy to ensure accurate concentration determination and labeling stoichiometry. The cells were washed twice with imaging buffer before imaging.

### FRET confocal microscopy

Fluorescence imaging was performed using confocal laser scanning microscopy (CLSM), on an Olympus IX81 confocal setup, with fixed acquisition settings within each dataset. Donor and acceptor fluorophores were sequentially excited, and emission signals were collected in separate channels to obtain directly excited donor (IDD), directly excited acceptor (IAA), and FRET-excited acceptor (IAD) intensities. Detailed acquisition settings are described in the SI Appendix.

### Image analysis and fluorescence quantification

Membrane-associated fluorescence intensities were quantified using an automated analysis pipeline based on line-scan extraction perpendicular to the plasma membrane. This approach enabled the acquisition of large datasets of spatially resolved fluorescence measurements across individual cells. Detailed image processing, segmentation, and line-scan analysis procedures are described in the SI Appendix.

### Correction of fluorescence signals

To account for donor bleed-through and direct acceptor excitation, fluorescence signals were corrected using experimentally determined linear coefficients derived from donor-only and acceptor-only conditions (Fig. S2). Corrected FRET signals were used to calculate Eapp,se for each measurement point. Full derivation of correction procedures is provided in the SI Appendix (Eqs. S1–S4).

### Calculation of apparent FRET efficiency

Apparent sensitized emission FRET efficiency (Eapp,se) between Alexa Fluor® 546 and Alexa Fluor® 647 was calculated using an approach described by Meyer *et al* (34):

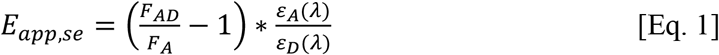

Where F_AD_ corresponds to the sum of acceptor emission when excited by the donor and the directly excited acceptor emission (IAA+IAD_corr_; SI Appendix), F_A_ corresponds to the directly excited acceptor emission (IAA), and *ε*_*D*_*(λ)*/*ε*_*A*_*(λ)* is the respective molar extinction ratio (0.172 in the case of Alexa Fluor® 546 and Alexa Fluor® 647).

For each intensity bin, the mean apparent FRET efficiency (Eapp,se), standard error of the mean (SEM), and number of observations (n) were calculated. These bin-averaged values were used for inter-experimental averaging and subsequent nonlinear regression analysis.

### Quantitative analysis of receptor oligomerization

Receptor oligomerization was quantified using two complementary analytical strategies. In Strategy 1, Eapp,se was analyzed as a function of receptor density using a monomer–dimer equilibrium model defined by an apparent dissociation constant (K_d_):

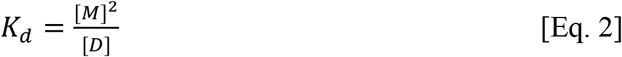

where [M] and [D] denote the concentrations of monomers and dimers, respectively. Total receptor concentration is given by:

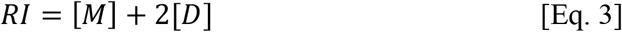

From this model, monomer and dimer fractions were derived to calculate the apparent oligomerization number:

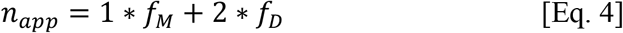

Under this definition, n_app_ ranges from 1 (fully monomeric) to 2 (fully dimeric). Full model equations and fitting procedures are provided in the SI Appendix (Eqs. S6–S12).

The model selection was based on goodness-of-fit metrics and model parsimony. Several models were evaluated, including a one-site specific binding model, a Hill equation (to account for potential cooperativity), and multi-exponential models. Hill fits were tested as an empirical alternative (Fig. S7) and yielded similar overall trends but did not improve the quality of the fits, when introducing an additional free parameter. Multi-exponential models were also explored but did not provide a consistent improvement in goodness-of-fit and were therefore not retained. The one-site binding model provided the most robust and parsimonious description of the data across datasets and was therefore used for all quantitative analyses presented in this study.

In Strategy 2, Eapp,se was analyzed as a function of donor fraction (x_D_) to independently estimate n_app_ and the intrinsic FRET efficiency (Etrue), based on established theoretical frameworks (34):

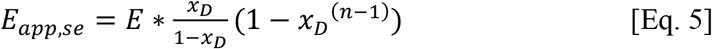

where E corresponds to the intrinsic pairwise FRET efficiency Etrue and n represents the apparent oligomeric state. Full model equations and fitting procedures are provided in the SI Appendix (Eq. S13).

### Statistical analysis and uncertainty estimation

All experiments were performed with multiple independent biological replicates. Data are presented as mean ± SEM unless otherwise stated. Parameter uncertainty was estimated using Monte Carlo simulations incorporating experimentally determined noise levels. Detailed statistical analysis and uncertainty propagation are described in the SI Appendix.

## Acknowledgments

The authors thank Nikos Hatzakis for accessing the Olympus IX81 confocal microscope (UCPH, DK). Plasmids were kindly provided by Martin Gustafsson (ST-CXCR4), Alexander S. Hauser (5-HT2AR), and Jonathan A. Javitch (ST-mGluR2 and ST-Δ2Δ). This work was supported by the Novo Nordisk Foundation grants NNF 20OC0064565 (K.L.M., M.M.R, P.M.B.) and NNF23OC008432 (K.L.M., M.M.R, P.M.B., S.M), the Danish Council for Strategic Research (1311-00002B) (K.L.M), and the Sino-Danish Center for Education and Research (K.L.M).

## Abbreviations

5-HT2AR: 5-hydroxytryptamine receptor 2A
AT1R: angiotensin II type 1 receptor
β1AR: beta-1 adrenergic receptor
β2AR: beta-2 adrenergic receptor
BRET: Bioluminescence Resonance Energy Transfer
CB1R: cannabinoid receptor 1
CXCR4: C-X-C motif chemokine receptor 4
Eapp,se: apparent sensitized-emission FRET efficiency
FRET: Förster Resonance Energy Transfer
GPCR: G protein–coupled receptor
IAA: directly excited acceptor intensity
IAD: FRET-excited acceptor intensity
IDD: directly excited donor intensity
K_d_: apparent homodimerization constant
MAS: MATLAB Automatic Cell Analysis Script
mGluR2: metabotropic glutamate receptor 2
n_app_: apparent oligomerization number
ST: SNAPtag
x_D_: donor-acceptor ratio

## Author contributions

C.D. and K.L.M. wrote the manuscript, with contributions from all the authors. C.D. and K.L.M. designed the project. C.D., N.B., E.P., M.M. and G.V.S.V.R. performed confocal imaging experiments, FRET measurements, and image acquisition. A.D. developed and performed automated image analysis with contributions from N.B. and J.H.. C.D. performed quantitative FRET analysis, and statistical analysis with contributions from A.D., N.B. and K.L.M.. C.D., K.L.M. discussed the experimental findings and interpretation of results. C.D. and K.L.M. supervised the project. with P.M.B., M.M.M., M.M.R. and S.M.

## Declaration of generative AI and AI-assisted technologies in the writing process

ChatGPT (OpenAI) and Copilot (Microsoft) were used for language editing, including improvements to clarity, grammar, and consistency. All AI-assisted text and code were reviewed and edited by the authors, who take full responsibility for the content of the manuscript

## Data availability / Code availability statements

Source data generated and analyzed that support the findings of this study will be shared by the lead contact upon request. Any additional information required to reanalyze the data reported in this paper is available from the lead contact upon request.

## Supporting Text

### FRET confocal microscopy

Confocal laser scanning microscopy (CLSM) was performed on an Olympus IX81 confocal setup using a UPLSAPO 100x oil immersion objective with a 1.4 numerical aperture. Alexa Fluor® 546 was excited by a 559 nm laser (25% laser power). Emission was captured in the range of 600-630 nm. Alexa Fluor® 647 was excited by a 635 nm laser (20% laser power). Emission was captured in the range of 670-770 nm. Detector gain and laser power were kept constant across experiments within each dataset (PMT settings: donor channel HV = 870, gain = 3x; acceptor channel HV = 722, gain = 2x). Images were acquired at a resolution of 1600 × 1600 pixels with identical acquisition settings for all conditions within an experiment (pixel dwell time: 2.0 µs/pixel).

### Automated membrane line-scan analysis

Membrane-associated fluorescence was quantified using a custom MATLAB script. Prior to analysis, confocal images were processed in FIJI to generate two combined image sets: (i) IAA_IAD, containing the directly excited acceptor (IAA) and the FRET-excited acceptor (IAD), and (ii) IAA_IDD, containing the directly excited acceptor (IAA) and the directly excited donor (IDD). Each dataset was analyzed independently with the custom MATLAB script using the two-membrane-channel mode.

Cell segmentation was performed on the receptor channel using a watershed-based regional minima algorithm. Images were pre-filtered using a median filter (radius = 3 pixels) to reduce high-frequency noise, followed by H-minima suppression with a maximum noise threshold of 20 to reduce over-segmentation. Objects smaller than 1,000 pixels or larger than 300,000 pixels were excluded from analysis.

Perpendicular line scans (25 pixels in length) were automatically drawn across membrane segments of segmented cells. For each line scan, intensity profiles were fitted with a Gaussian function to extract membrane-associated signal as the integrated area under the Gaussian, while the offset was defined as background intensity. Line scans were accepted only if the full width at half maximum (FWHM) was < 15 pixels and the goodness-of-fit satisfied R^2^ ≥ 0.9. Line scans intersecting nuclei were excluded using a nucleus intensity threshold of 5,000 arbitrary units.

The two MATLAB analyses yielded IAA, IAD, and IDD intensity values, which were subsequently merged at the line-scan level for downstream correction of spectral bleed-through and calculation of apparent FRET efficiency.

### Data consolidation

Following independent analyses of the IAA_IAD and IAA_IDD image sets, output .csv files containing line-scan intensities were merged at the individual line-scan level. For each line scan, directly excited acceptor intensity (IAA), FRET-excited acceptor intensity (IAD), and directly excited donor intensity (IDD) were combined into a single dataset for downstream quantitative analysis.

### Fluorescence intensities correction

To ensure that apparent FRET efficiency measurements (Eapp,se) reflect receptor-specific oligomerization rather than optical cross-talk, correction equations were established to remove nonspecific intermolecular FRET arising from direct acceptor excitation and donor bleed-through (75). At donor fraction x_D_ = 0 (acceptor only), the acceptor channel signal upon donor excitation originates exclusively from direct cross-excitation of the acceptor fluorophore. Under these conditions, the acceptor signal (IAD) scales linearly with acceptor emission upon acceptor excitation (IAA), allowing estimation of the direct excitation contribution (Fig. S2A). Because measurements were acquired over an extended time period, sight variations in instrument performance required the use of dataset-specific correction equations. Similarly, measurements performed at x_D_ = 1 (donor only) isolate donor emission bleed-through into the acceptor channel. In this configuration, IAD varies linearly with donor emission upon donor excitation (IDD), providing an estimate of donor bleed-through (Fig. S2B).

Based on these control conditions, linear correction coefficients were determined experimentally. The donor bleed-through coefficient (*a*) was determined from donor-only samples (x_D_ = 1) as:

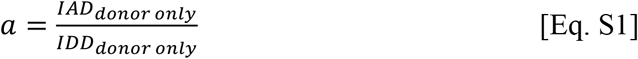

where IAD_donor only_ represents the signal detected in the acceptor channel upon donor excitation and IDD_donor only_ corresponds to the directly excited donor intensity.

The direct acceptor excitation coefficient (*d*) was determined from acceptor-only samples (x_D_ = 0) as:

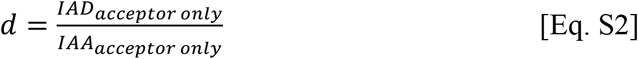

Where IAD_acceptor only_ represents the acceptor signal detected under donor excitation and IAA_acceptor only_ corresponds to the directly excited acceptor intensity.

Corrected FRET intensity was calculated for each line scan according to:

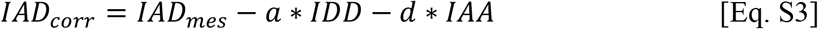

where IAD_mes_ is the measured acceptor-channel signal under donor excitation.

For live-cell experiments performed under the optical configuration used in this study, correction factors were determined empirically for each acquisition period. For all datasets except β1AR, the correction equation was:

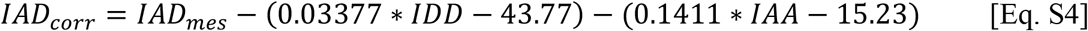

Although identical acquisition settings and optical configuration were used, correction factors for the β1AR dataset were determined independently (Fig. S2A-B), as these parameters can vary over time due to changes in laser performance and detector sensitivity.

### Intensity normalization and binning strategy

Membrane receptor density (RI) was estimated using the following equation:

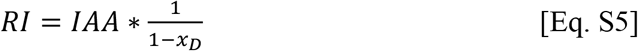

### Nonlinear regression and oligomerization modeling

#### Strategy 1 – density-dependent saturation analysis

To estimate the apparent dissociation constant (K_d_) and maximal apparent FRET efficiency (Eapp_max_), nonlinear regression was performed at donor molar fraction x_D_ = 0.5. Bin-averaged Eapp,se values were fitted as a function of membrane receptor density (RI) using a one-site specific binding model on GraphPad Prism (GraphPad Software). Best-fit parameters were obtained by least-squares regression and reported as K_d_ and Eapp_max_.

Assuming a monomer–dimer equilibrium (2M ⇌ D), the apparent dissociation constant is defined as:

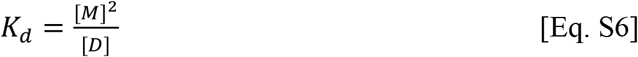

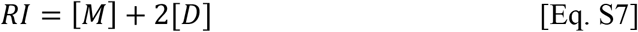

For a given total receptor density (RI), monomer concentration was calculated analytically as:

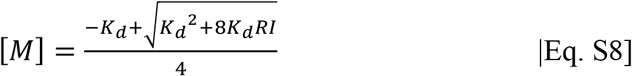

Dimer concentration was then obtained from:

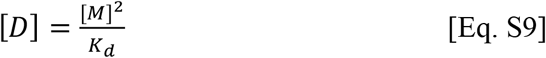

The fraction of receptors in monomeric and dimeric states was calculated as:

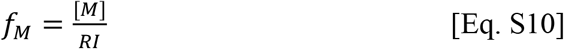

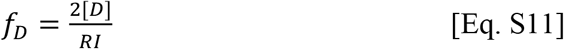

such that *f*_*M*_+*f*_*D*_ = 1.

The apparent oligomeric state was defined as the fraction-weighted average oligomer size:

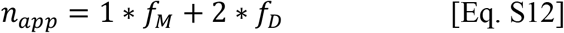

Under this definition, n_app_ ranges from 1 (fully monomeric) to 2 (fully dimeric).

#### Strategy 2 – x_D_-dependent analysis

In addition to density-dependent saturation analysis (Strategy 1), the intrinsic pairwise FRET efficiency (Etrue) and apparent oligomeric state were evaluated from the dependence of apparent sensitized-emission FRET efficiency (Eapp,se) on the x_D_, using Equation 8 from Meyer *et al*. (34).

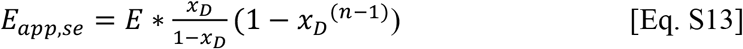

where E corresponds to the intrinsic pairwise FRET efficiency Etrue and n represents the apparent oligomeric state.

Because this approach relies on a limited number of x_D_ conditions compared to the large number of data points available in density-dependent saturation curves, parameter estimation from this fit is inherently less precise than Strategy 1. Therefore, whenever a reliable K_d_ and n_app_ could be obtained from Strategy 1, the value of n_app_ determined from the density-dependent analysis was fixed in [Equation S13] to calculate Etrue. The Eapp,se = f(x_D_) relationship was then refitted using Etrue as the primary parameter.

For receptors where density-dependent analysis did not yield a reliable K_d_, both Etrue and n were estimated from the x_D_-dependent fit. In cases where neither strategy provided a stable estimate of Etrue, the intrinsic FRET efficiency determined for a structurally related receptor within the same GPCR class (class A or class C) was used in Equation 8 to estimate n_app_.

### Statistical analysis

All experiments were performed with a minimum of three independent biological replicates per receptor. For each experiment, line-scan data were first averaged within intensity-based bins, and bin means were subsequently averaged across independent experiments. Data are presented as mean ± SEM unless otherwise stated. Nonlinear regression analyses were performed using GraphPad Prism (GraphPad Software).

The intrinsic experimental noise of the method was estimated from the dispersion of Eapp,se values measured for the monomeric control receptor Δ2Δ, which is not expected to oligomerize. The root mean square error (RMSE) of the Eapp,se distribution across the explored receptor density range was calculated and yielded a value of 0.00188. Assuming normally distributed measurement errors, the practical detection limit of the method was defined as three times the RMSE (3 × RMSE ≈ 0.0056), corresponding to the minimal variation in apparent FRET efficiency that can be reliably distinguished from experimental noise.

In addition to this intrinsic noise estimate, the sensitivity of the method to detect density-dependent changes in FRET was assessed from the dispersion of slopes obtained by linear regression of Eapp,se as a function of receptor density for receptors expected to exhibit density-independent oligomerization states (Δ2Δ and mGluR2). The standard deviation of these slopes across independent experiments was used to estimate the minimal detectable density-dependent change, assuming normally distributed noise.

In Strategy 1, Eapp,se values obtained at a donor molar fraction x_D_ = 0.5 were fitted as a function of receptor intensity using a monomer–dimer equilibrium model to estimate the dissociation constant (K_d_) and the maximum apparent FRET efficiency (Eapp_max_). From these parameters, total receptor concentration (RT) and the fractions of monomers and dimers were calculated assuming a two-state equilibrium, and the apparent oligomeric state (n_app_) was derived accordingly.

Uncertainty in fitted parameters was estimated using Monte Carlo simulations. The best-fit model was used as the underlying ground truth, and Gaussian noise corresponding to the experimentally determined RMSE was added to generate 1000 simulated datasets. Each dataset was refitted using the same model to obtain a distribution of K_d_, from which corresponding values of n_app_, receptor populations, and intrinsic FRET efficiency (E) were derived.

In Strategy 2, the relationship between Eapp,se and donor molar fraction (x_D_) was analyzed using the Equation 5 to estimate n_app_. When a reliable estimate of the intrinsic FRET efficiency (E) was obtained from Strategy 1, this value was fixed during fitting to reduce parameter degeneracy. To propagate uncertainty from Strategy 1 into Strategy 2, a Monte Carlo approach was implemented. Specifically, 1000 values of E derived from Strategy 1 simulations were used as fixed inputs, and for each value, the Eapp,se versus x_D_ relationship was refitted to estimate a corresponding n_app_. Experimental variability in Eapp,se was incorporated by resampling data points according to their SEM. The resulting distributions of n_app_ were then converted into monomer and dimer fractions, and into [M] and [D], assuming a monomer–dimer equilibrium. Final estimates and confidence intervals were obtained from the empirical distributions (median and 2.5–97.5th percentiles).

## Supporting Figures

**Figure S1:**
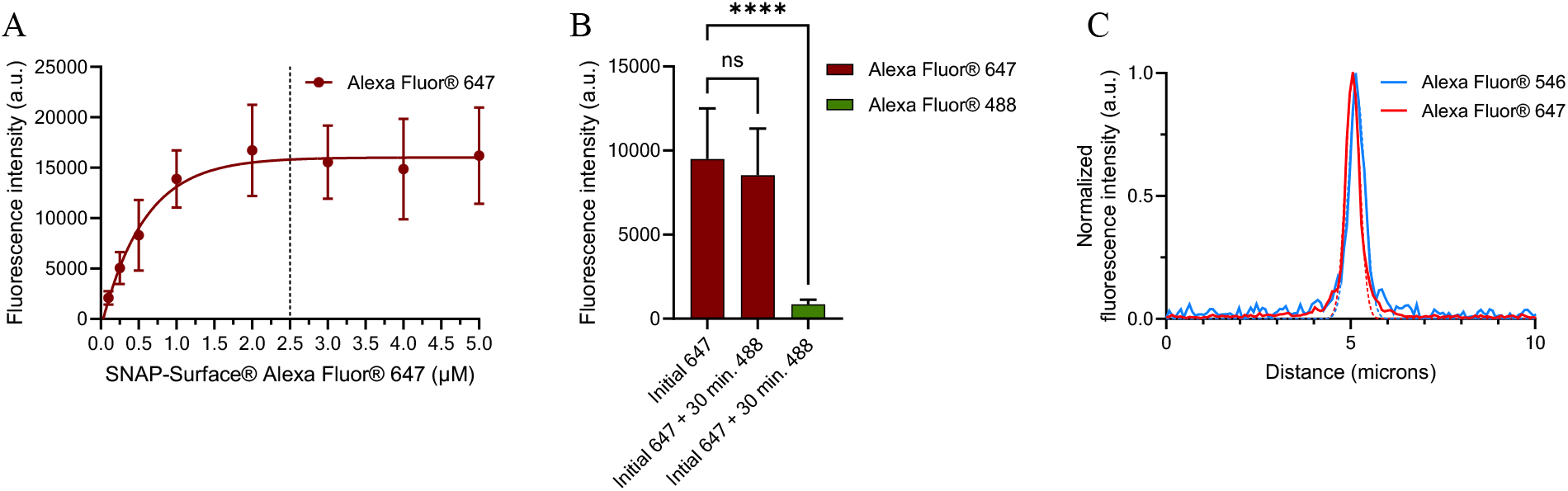
Validation of SNAP labeling conditions in iSNAP-β1AR cells. (A) Fluorescence intensity as a function of SNAP-Surface® Alexa Fluor 647 concentration in iSNAP-β1AR cells. Cells were incubated for 30 min with increasing concentrations of SNAP-Surface® Alexa Fluor 647. Fluorescence intensity reached a plateau at ∼2.5 µM, indicating saturation of receptor labeling. Data is shown as mean ± SD. (B) Assessment of labeling saturation. Cells were first labeled with SNAP-Surface® Alexa Fluor 647 (2.5 µM, 30 min), washed, and subsequently incubated with SNAP-Surface® Alexa Fluor 488 (2.5 µM, 30 min). Minimal Alexa Fluor 488 signal was detected (third column) compared with the initial Alexa Fluor 647 labeling (second column), indicating that SNAP sites were saturated during the first labeling step. Statistical analysis was performed using an ordinary one-way ANOVA. ns, not significant; ****P < 0.0001. (C) Fluorescence intensity profiles obtained from a 10-µm line scan across the plasma membrane of iSNAP-β1AR cells labeled with SNAP-Surface® Alexa Fluor 546 or Alexa Fluor 647. Normalized fluorescence intensity plotted as a function of distance shows similar peak amplitudes and spatial distributions for both fluorophores, indicating comparable labeling efficiency under the experimental conditions.

**Figure S2:**
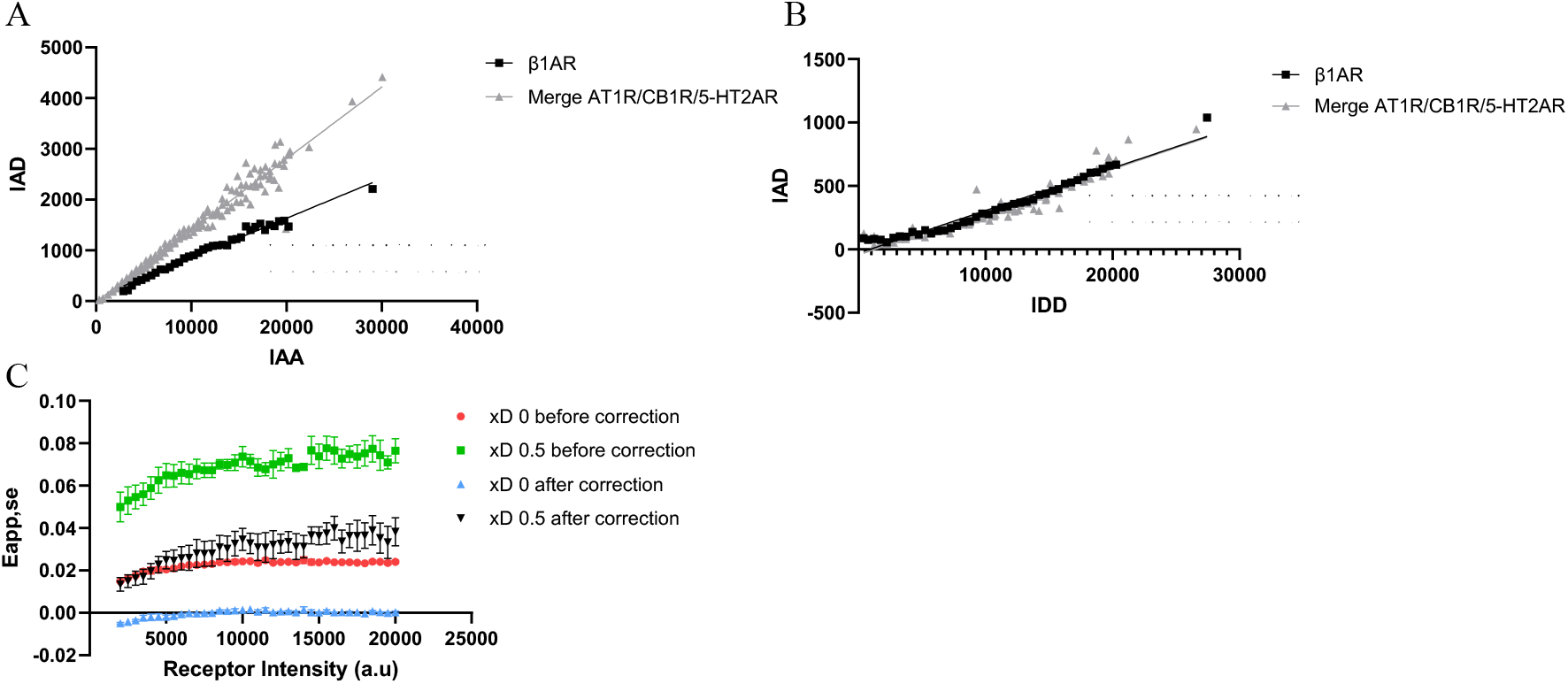
Validation of the IAD correction used to remove nonspecific FRET contributions. (A) Acceptor emission upon donor excitation (IAD) plotted as a function of acceptor emission upon acceptor excitation (IAA) at donor fraction x_D_ = 0, where no donor– acceptor FRET occurs. Linear relationships obtained for the β1AR dataset and for a merged dataset of AT1R, CB1R, and 5-HT2AR receptors are shown together with the corresponding regression fits used for correction of direct acceptor excitation. (B) IAD plotted as a function of donor emission upon donor excitation (IDD) at x_D_ = 1, where the acceptor channel signal originates exclusively from donor emission bleed-through. Linear regression fits used for bleed-through correction are shown. (C) Eapp,se plotted as a function of receptor surface density in HEK293 cells expressing ST–AT1R at x_D_ = 0 and x_D_ = 0.5, before and after application of the IAD correction. Data corresponds to n=2 independent experiments for x_D_ = 0 and n = 4 for x_D_ = 0.5.

**Figure S3:**
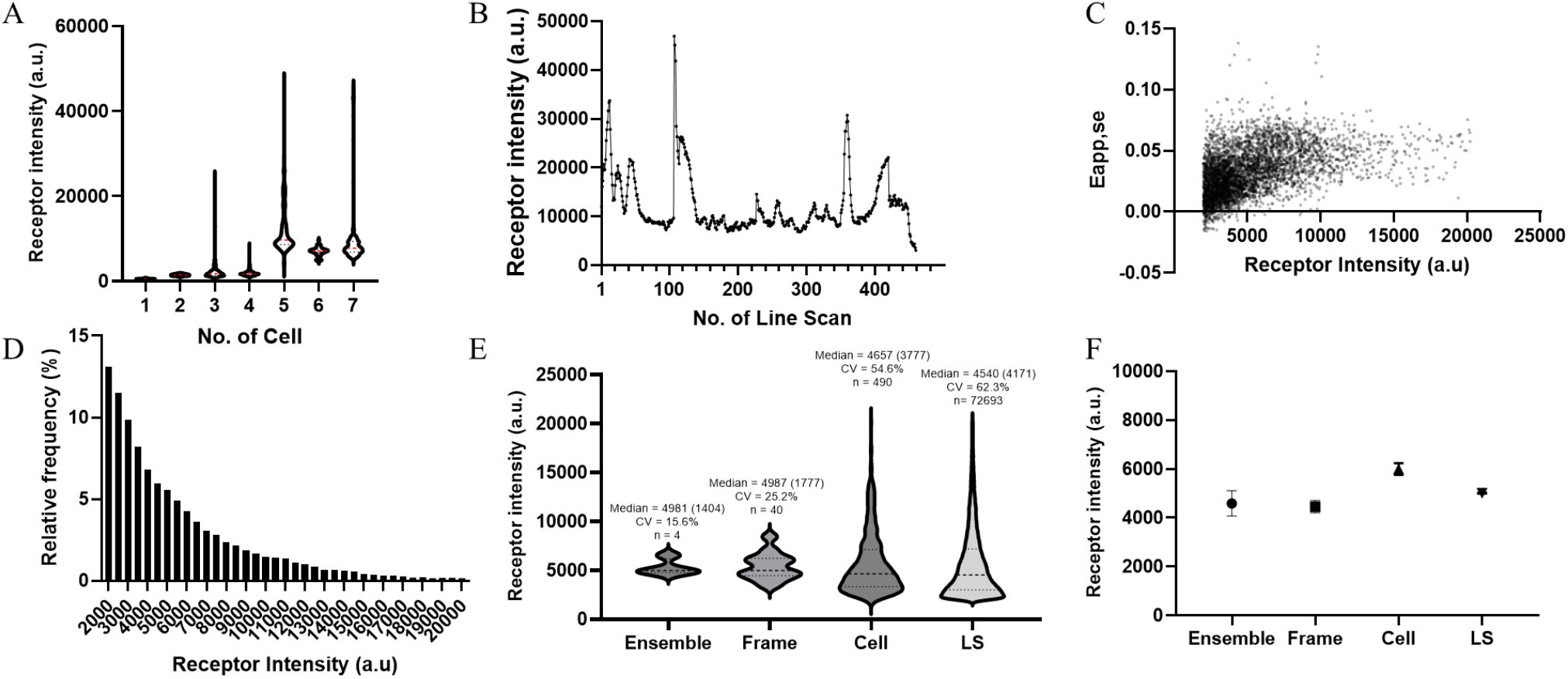
Characterization of receptor density heterogeneity and construction of large datasets for quantitative FRET analysis. (A-B) Representative example of receptor heterogeneity on one frame by the distribution of the receptor fluorescence intensities measured across individual HEK293 cells transiently expressing ST-AT1R (A) and along the plasma membrane of cell No. 5 (B), measured at a x_D_ of 0.5. (C) Eapp,se plotted as a function of receptor intensity obtained from individual line scan (LS). For visualization purposes, a random subset of 5,000 measurements from the full dataset is shown. (D) Number of line scan measurements obtained in HEK293 cell transiently expressing ST-AT1R, measured at a x_D_ of 0.5 (4 independent experiments). The histogram values are normalized to percentage of total counts. (E) Violin plots showing the distribution of receptor intensities calculated at different levels of data aggregation: the experiment level (Ensemble; one value per independent experiment, n = 4), individual image frames (Frame; n = 40), individual cells (Cell; n = 490), and individual line scan (LS; n = 72693). Median values (with IQR in parentheses) and coefficients of variation (CV) are indicated above each distribution and mean ± SEM are shown in (F).

**Figure S4:**
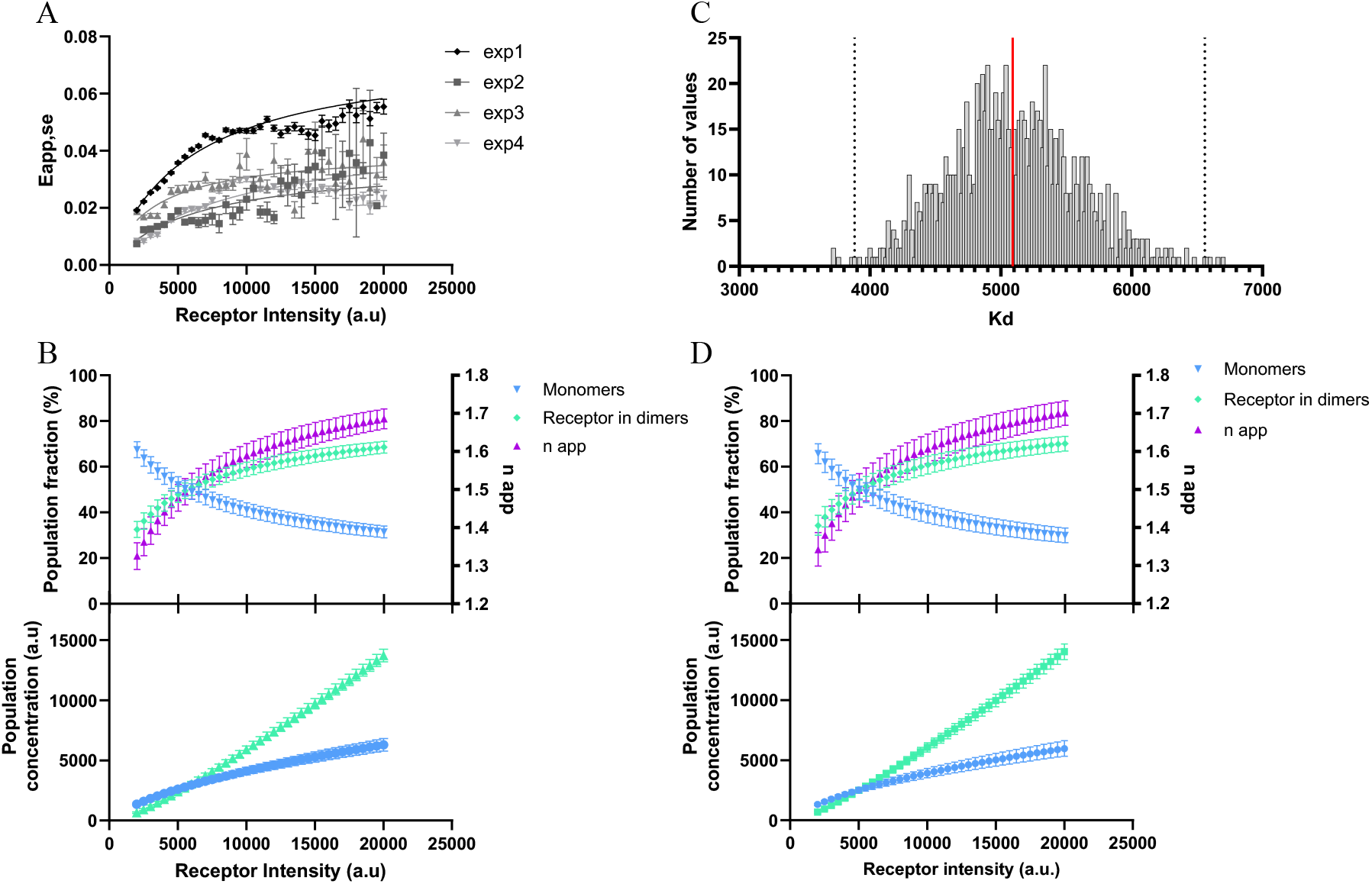
Experimental variability and uncertainty propagation in the estimation of AT1R oligomerization. (A) Eapp,se plotted as a function of receptor surface density (a.u) in HEK293 cells transiently expressing ST-AT1R, measured at a x_D_ of 0.5. Each dataset corresponds to the mean ± SEM obtained from 4 independent experiments. Data were fitted independently using the one-site specific binding model (GraphPad Prism), to estimate the K_d_. (B) Fractions of monomeric and dimeric AT1R populations are estimated as a function of receptor surface density using the K_d_ values obtained in (A). Monomer and dimer populations are expressed as percentages (left y-axis), while the corresponding n app is shown on the right y-axis. The corresponding concentrations of monomeric and dimeric receptor species as a function of receptor surface density are shown directly below. Each point represents the mean ± SEM obtained from n = 4 independent experiments for AT1R. (C) Distribution of K_d_ values obtained from Monte Carlo simulations (n = 1000) incorporating experimental noise. The red line indicates the mean K_d,_ and dashed lines indicate the 99% confidence interval. Fractions of monomeric and dimeric AT1R populations are estimated as a function of receptor surface density using the apparent K_d_ obtained in C. Monomer and dimer populations are expressed as percentages (left y-axis), while the corresponding n app is shown on the right y-axis. The corresponding concentrations of monomeric and dimeric receptor species as a function of receptor surface density are shown directly below. Each point represents the mean ± 99% CI obtained from the 1000 Monte-Carlo simulations.

**Figure S5:**
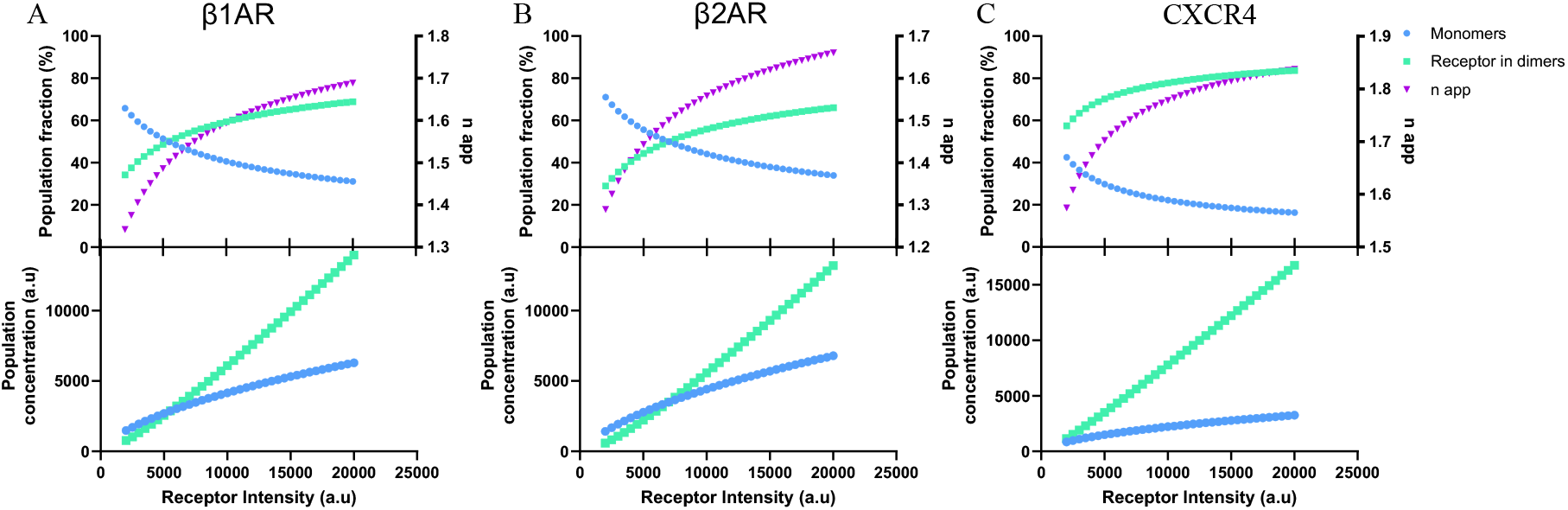
Predicted receptor populations derived from fitted K_d_ values for class A GPCRs. (A-C) Using the K_d_ value derived from the one-site specific binding model fit in Figure 3, the fractions of monomeric and dimeric receptors were estimated and plotted as percentages as a function of receptor surface density (left y-axis), while the corresponding apparent oligomerization number (n_app_) is shown on the right y-axis. The corresponding concentrations of monomeric and dimeric receptor species as a function of receptor surface density are shown directly below. Calculations were performed for β1AR (A), β2AR (B), and CXCR4 (C) using the K_d_ values obtained from the one-site specific binding model fits. The reported K_d_ values were derived from at least three independent experiments.

**Figure S6:**
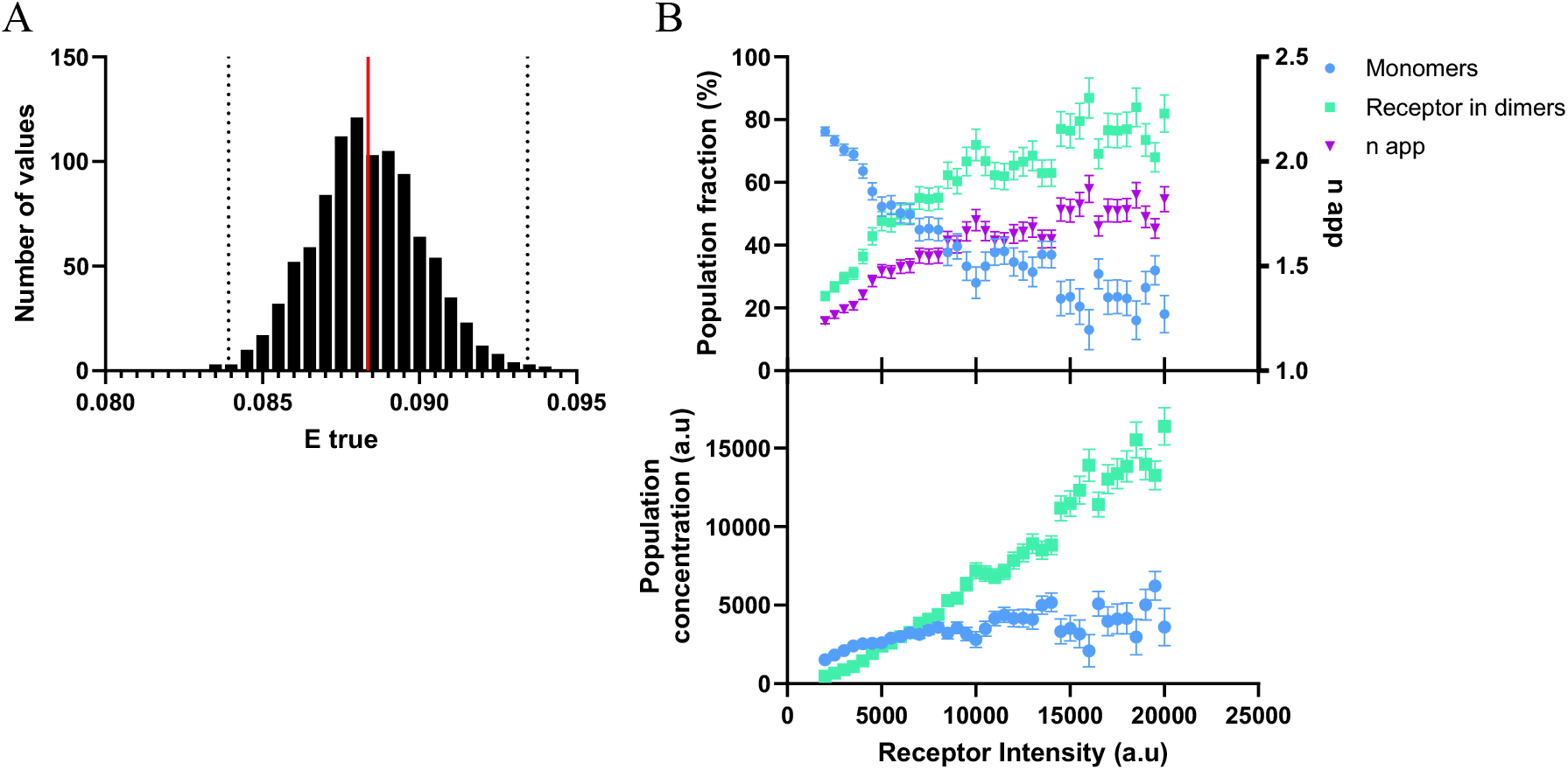
Uncertainty propagation in the estimation of AT1R oligomerization with the second Strategy. (A) Distribution of intrinsic FRET efficiencies (Etrue) obtained from Monte Carlo simulations. A total of 1000 dissociation constants K_d_ were randomly sampled according to the uncertainty estimated in Strategy 1. For each simulated K_d_, the corresponding apparent oligomeric state (n_app_) was calculated, and the Etrue was derived. The histogram shows the resulting distribution of the 1000 simulated Etrue values. The red line indicates the mean Etrue, and dashed lines indicate the 99% confidence interval. (B) Propagation of the simulated Etrue values into Strategy 2. Each Etrue value was fixed in Eq. 5 to estimate the n app from the relationship between Eapp,se and x_D_. The relationship used for this analysis corresponds to the AT1R dataset. The resulting n_app_ values are expressed as percentages (right y-axis), while the corresponding monomer and dimer populations are shown on the left y-axis. The corresponding concentrations of monomeric and dimeric receptor species as a function of receptor surface density are shown directly below. Each point represents the mean ± 99% CI obtained from the 1000 Monte-Carlo simulations.

**Figure S7:**
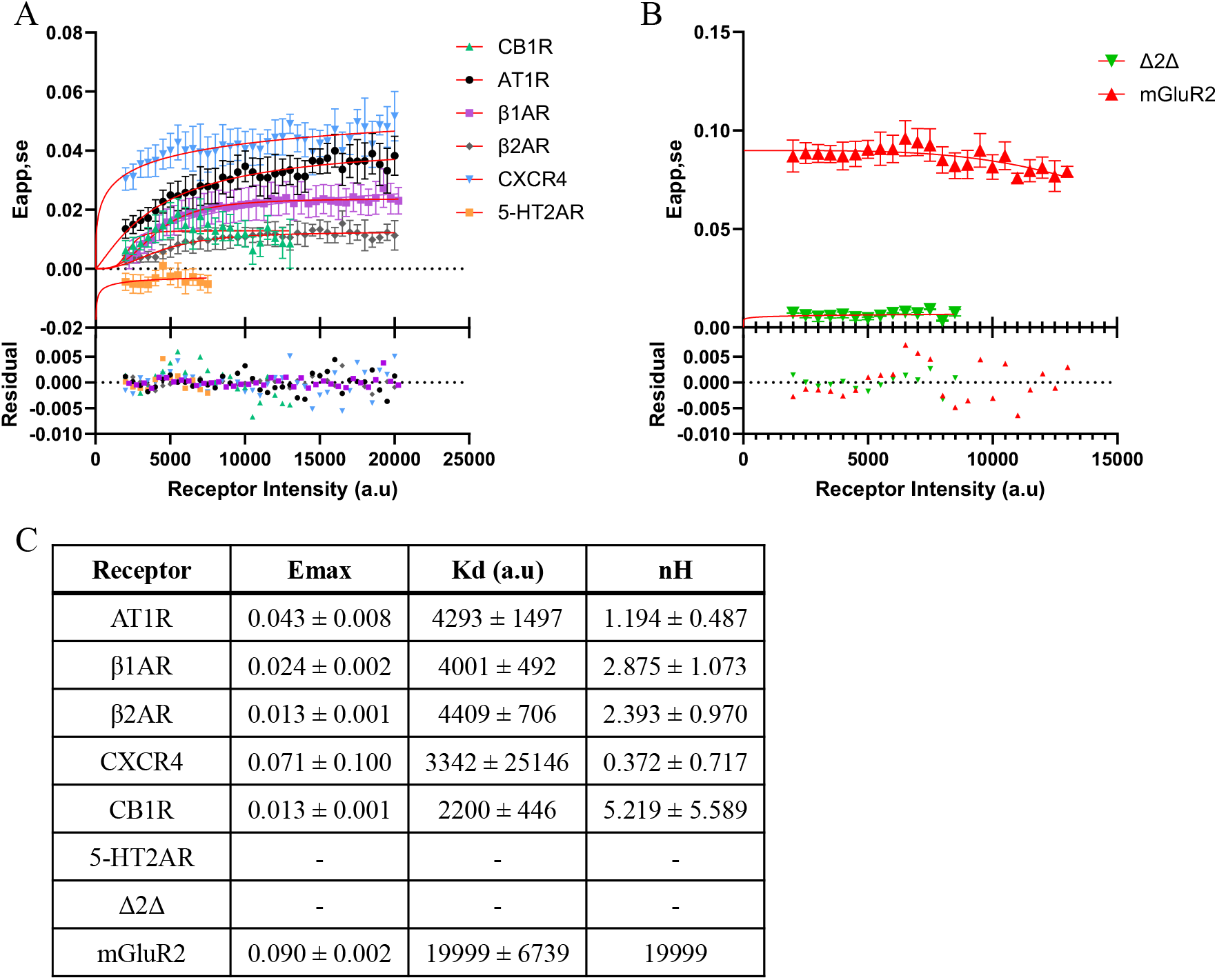
Comparative analysis of density-dependent oligomerization across class A GPCRs and controls. (A-B) Eapp,se plotted as a function of receptor surface density (a.u.) in HEK293 cells transiently expressing SNAP-tagged class A GPCRs, including AT1R, β1AR, β2AR, CXCR4, CB1R, and 5-HT2AR (A), or mGluR2 and Δ2Δ (B). Symbols represent mean ± SEM of experimental data, and solid lines indicate nonlinear regression using the specific binding model with Hill slope (GraphPad Prism). All receptors were analyzed using a minimum of n = 3 independent experiments except Δ2Δ (n = 2). (C) Summary of apparent dissociation constants (K_d_), Hill coefficients (nH), and maximal FRET efficiencies (Emax) derived from the fits applied to the data in A and B. Negative or nonconvergent K_d_ values indicate the absence of detectable density-dependent population changes within the explored expression range, consistent with predominantly monomeric or dimeric receptor populations.

