## supplementary material for "A quantitative imaging framework reveals density-dependent GPCR oligomerization and organization in living cells"

Based on these control conditions, linear correction coefficients were determined experimentally. The donor bleed-through coefficient (*a*) was determined from donor-only samples ($x_D = 1$) as:

$$a = \frac{IAD_{donor\ only}}{IDD_{donor\ only}} \qquad \text{[Eq. S1]}$$

where $IAD_{donor\ only}$ represents the signal detected in the acceptor channel upon donor excitation and $IDD_{donor\ only}$ corresponds to the directly excited donor intensity.

The direct acceptor excitation coefficient (*d*) was determined from acceptor-only samples ($x_D$ = 0) as:

$$d = \frac{IAD_{acceptor\ only}}{IAA_{acceptor\ only}} \qquad \text{[Eq. S2]}$$

Where $IAD_{acceptor\ only}$ represents the acceptor signal detected under donor excitation and $IAA_{acceptor\ only}$ corresponds to the directly excited acceptor intensity.

Corrected FRET intensity was calculated for each line scan according to:

$$IAD_{corr} = IAD_{mes} - a * IDD - d * IAA \qquad \text{[Eq. S3]}$$

where $IAD_{mes}$ is the measured acceptor-channel signal under donor excitation.

For live-cell experiments performed under the optical configuration used in this study, correction factors were determined empirically for each acquisition period. For all datasets except β1AR, the correction equation was:

$$IAD_{corr} = IAD_{mes} - (0.03377 * IDD - 43.77) - (0.1411 * IAA - 15.23) \qquad \text{[Eq. S4]}$$

Although identical acquisition settings and optical configuration were used, correction factors for the β1AR dataset were determined independently (Fig. S2A-B), as these parameters can vary over time due to changes in laser performance and detector sensitivity.

**Intensity normalization and binning strategy**

Membrane receptor density (RI) was estimated using the following equation:

$$RI = IAA * \frac{1}{1-x_D} \qquad \text{[Eq. S5]}$$

**Nonlinear regression and oligomerization modeling**

**Strategy 1 – density-dependent saturation analysis:**

To estimate the apparent dissociation constant ($K_d$) and maximal apparent FRET efficiency ($Eapp_{max}$), nonlinear regression was performed at donor molar fraction $x_D = 0.5$. Bin-averaged Eapp,se values were fitted as a function of membrane receptor density (RI) using a one-site specific binding model on GraphPad Prism (GraphPad Software). Best-fit parameters were obtained by least-squares regression and reported as $K_d$ and $Eapp_{max}$.

Assuming a monomer–dimer equilibrium (2M ⇌ D), the apparent dissociation constant is defined as:

$$K_d = \frac{[M]^2}{[D]} \qquad \text{[Eq. S6]}$$

where [M] and [D] denote the concentrations of monomers and dimers, respectively. Total receptor concentration is given by:

$$RI = [M] + 2[D] \qquad \text{[Eq. S7]}$$

For a given total receptor density (RI), monomer concentration was calculated analytically as:

$$[M] = \frac{-K_d + \sqrt{K_d{}^2 + 8K_d RI}}{4} \qquad \text{|Eq. S8]}$$

Dimer concentration was then obtained from:

$$[D] = \frac{[M]^2}{K_d} \qquad \text{[Eq. S9]}$$

The fraction of receptors in monomeric and dimeric states was calculated as:

$$f_M = \frac{[M]}{RI} \qquad \text{[Eq. S10]}$$

$$f_D = \frac{2[D]}{RI} \qquad \text{[Eq. S11]}$$

such that $f_M + f_D = 1$.

The apparent oligomeric state was defined as the fraction-weighted average oligomer size:

### Supporting Figures

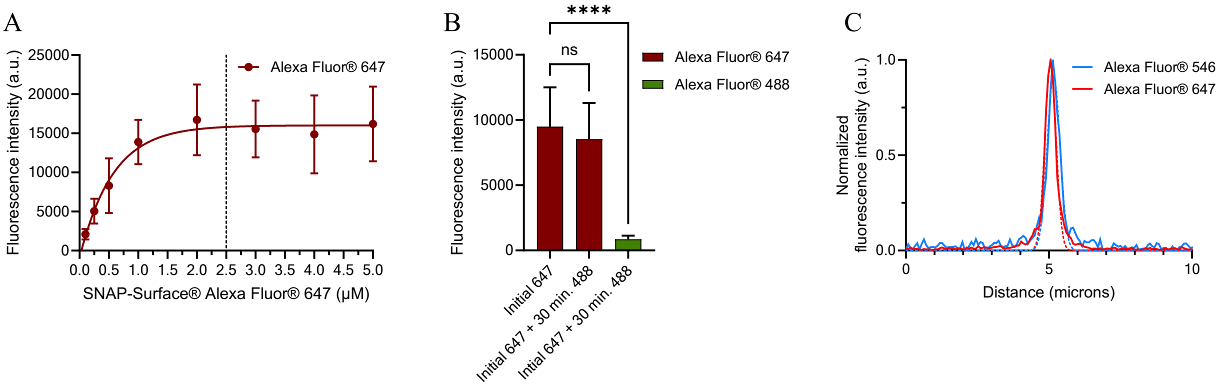


**Figure S1: Validation of SNAP labeling conditions in iSNAP-β1AR cells**. (A) Fluorescence intensity as a function of SNAP-Surface® Alexa Fluor 647 concentration in iSNAP-β1AR cells. Cells were incubated for 30 min with increasing concentrations of SNAP-Surface® Alexa Fluor 647. Fluorescence intensity reached a plateau at ~2.5 µM, indicating saturation of receptor labeling. Data is shown as mean ± SD. (B) Assessment of labeling saturation. Cells were first labeled with SNAP-Surface® Alexa Fluor 647 (2.5 µM, 30 min), washed, and subsequently incubated with SNAP-Surface® Alexa Fluor 488 (2.5 µM, 30 min). Minimal Alexa Fluor 488 signal was detected (third column) compared with the initial Alexa Fluor 647 labeling (second column), indicating that SNAP sites were saturated during the first labeling step. Statistical analysis was performed using an ordinary one-way ANOVA. ns, not significant; ****P < 0.0001. (C) Fluorescence intensity profiles obtained from a 10-µm line scan across the plasma membrane of iSNAP-β1AR cells labeled with SNAP-Surface® Alexa Fluor 546 or Alexa Fluor 647. Normalized fluorescence intensity plotted as a function of distance shows similar peak amplitudes and spatial distributions for both fluorophores, indicating comparable labeling efficiency under the experimental conditions.

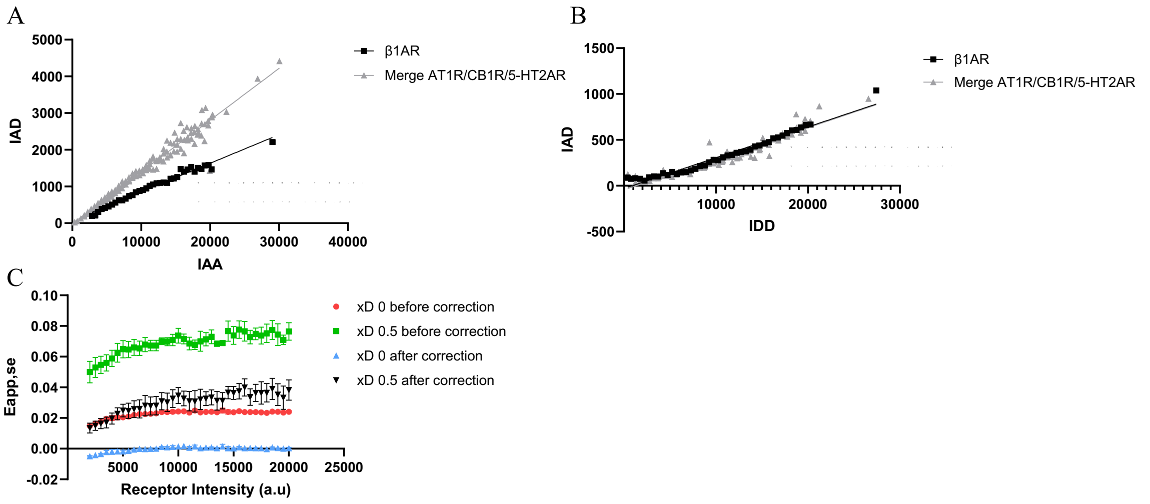


**Figure S2: Validation of the IAD correction used to remove nonspecific FRET contributions.** (A) Acceptor emission upon donor excitation (IAD) plotted as a function of acceptor emission upon acceptor excitation (IAA) at donor fraction $x_D = 0$, where no donor–

acceptor FRET occurs. Linear relationships obtained for the β1AR dataset and for a merged dataset of AT1R, CB1R, and 5-HT2AR receptors are shown together with the corresponding regression fits used for correction of direct acceptor excitation. (B) IAD plotted as a function of donor emission upon donor excitation (IDD) at $x_D = 1$, where the acceptor channel signal originates exclusively from donor emission bleed-through. Linear regression fits used for bleed-through correction are shown. (C) Eapp,se plotted as a function of receptor surface density in HEK293 cells expressing ST–AT1R at $x_D = 0$ and $x_D = 0.5$, before and after application of the IAD correction. Data corresponds to n=2 independent experiments for $x_D = 0$ and n = 4 for $x_D = 0.5$.

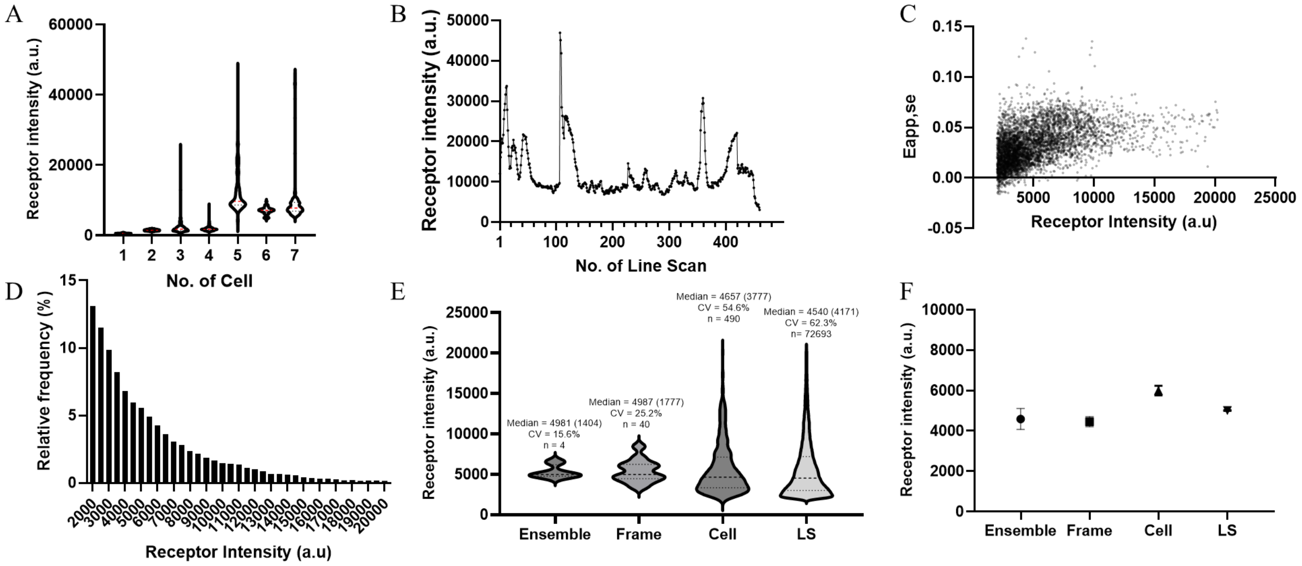


**Figure S3: Characterization of receptor density heterogeneity and construction of large datasets for quantitative FRET analysis.** (A-B) Representative example of receptor heterogeneity on one frame by the distribution of the receptor fluorescence intensities measured across individual HEK293 cells transiently expressing ST-AT1R (A) and along the plasma membrane of cell No. 5 (B), measured at a $x_D$ of 0.5. (C) Eapp,se plotted as a function of receptor intensity obtained from individual line scan (LS). For visualization purposes, a random subset of 5,000 measurements from the full dataset is shown. (D) Number of line scan measurements obtained in HEK293 cell transiently expressing ST-AT1R, measured at a $x_D$ of 0.5 (4 independent experiments). The histogram values are normalized to percentage of total counts. (E) Violin plots showing the distribution of receptor intensities calculated at different levels of data aggregation: the experiment level (Ensemble; one value per independent experiment, n = 4), individual image frames (Frame; n = 40), individual cells (Cell; n = 490), and individual line scan (LS; n = 72693). Median values (with IQR in parentheses) and coefficients of variation (CV) are indicated above each distribution and mean ± SEM are shown in (F).

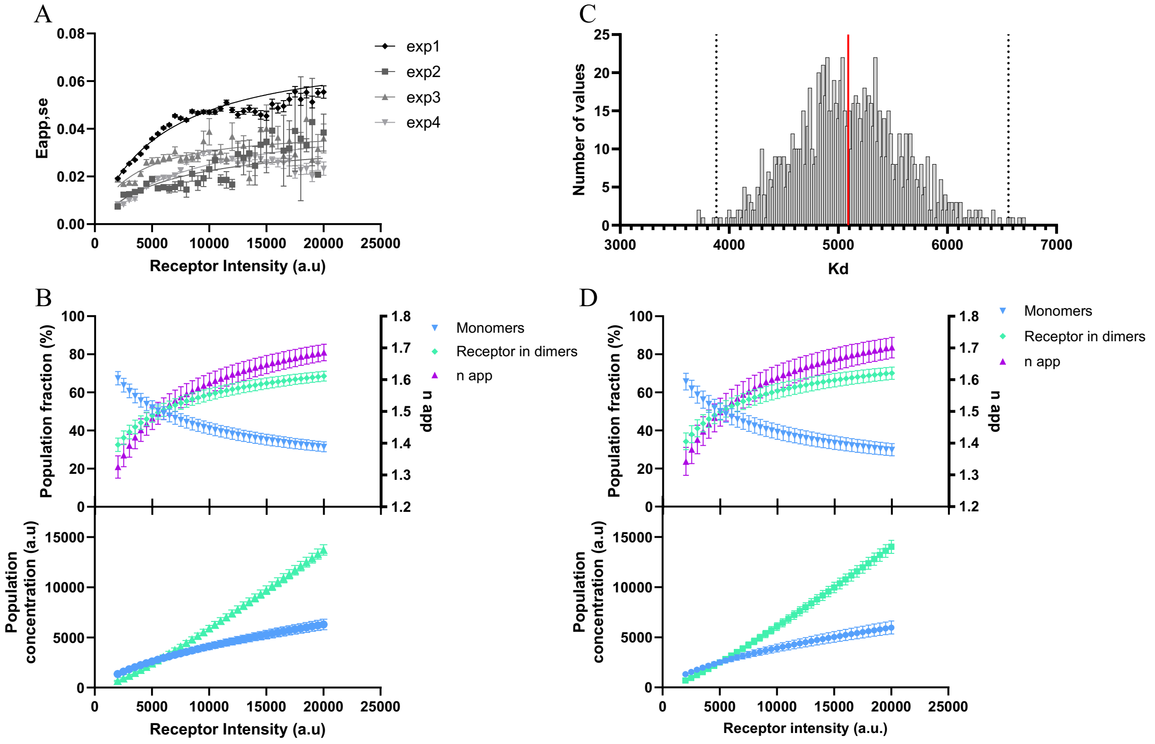


**Figure S4: Experimental variability and uncertainty propagation in the estimation of AT1R oligomerization.** (A) Eapp,se plotted as a function of receptor surface density (a.u) in HEK293 cells transiently expressing ST-AT1R, measured at a $x_D$ of 0.5. Each dataset corresponds to the mean ± SEM obtained from 4 independent experiments. Data were fitted independently using the one-site specific binding model (GraphPad Prism), to estimate the $K_d$. (B) Fractions of monomeric and dimeric AT1R populations are estimated as a function of receptor surface density using the $K_d$ values obtained in (A). Monomer and dimer populations are expressed as percentages (left y-axis), while the corresponding n app is shown on the right y-axis. The corresponding concentrations of monomeric and dimeric receptor species as a function of receptor surface density are shown directly below. Each point represents the mean ± SEM obtained from n = 4 independent experiments for AT1R. (C) Distribution of $K_d$ values obtained from Monte Carlo simulations (n = 1000) incorporating experimental noise. The red line indicates the mean $K_{d,}$ and dashed lines indicate the 99% confidence interval. (D) Fractions of monomeric and dimeric AT1R populations are estimated as a function of receptor surface density using the apparent $K_d$ obtained in C. Monomer and dimer populations are expressed as percentages (left y-axis), while the corresponding n app is shown on the right y-axis. The corresponding concentrations of monomeric and dimeric receptor species as a function of receptor surface density are shown directly below. Each point represents the mean ± 99% CI obtained from the 1000 Monte-Carlo simulations.

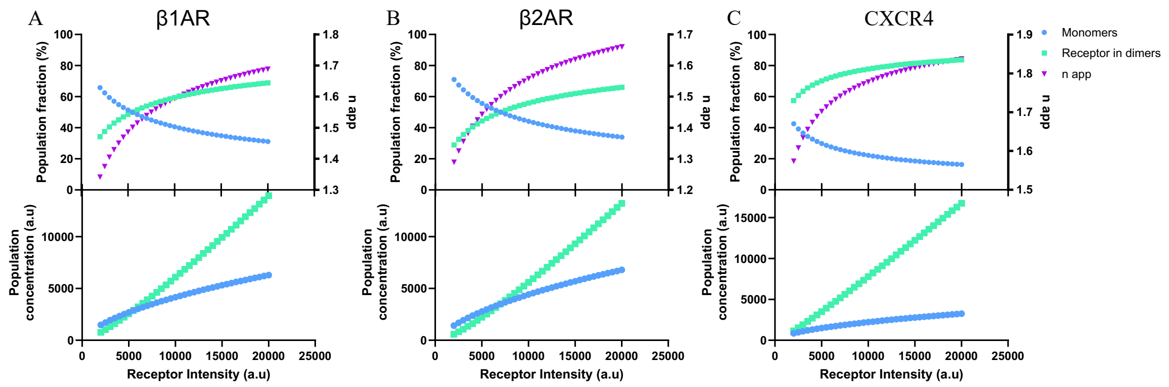


**Figure S5: Predicted receptor populations derived from fitted $K_d$ values for class A GPCRs.** (A-C) Using the $K_d$ value derived from the one-site specific binding model fit in Figure 3, the fractions of monomeric and dimeric receptors were estimated and plotted as percentages as a function of receptor surface density (left y-axis), while the corresponding apparent oligomerization number ($n_{app}$) is shown on the right y-axis. The corresponding concentrations of monomeric and dimeric receptor species as a function of receptor surface density are shown directly below. Calculations were performed for β1AR (A), β2AR (B), and CXCR4 (C) using the $K_d$ values obtained from the one-site specific binding model fits. The reported $K_d$ values were derived from at least three independent experiments.

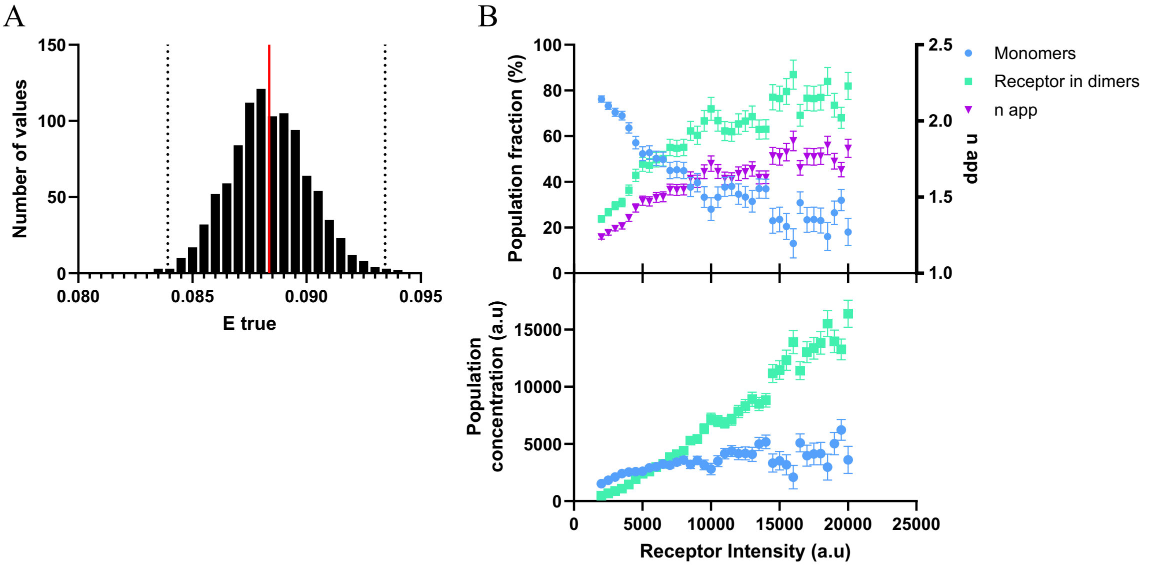


**Figure S6: Uncertainty propagation in the estimation of AT1R oligomerization with the second Strategy.** (A) Distribution of intrinsic FRET efficiencies (Etrue) obtained from Monte Carlo simulations. A total of 1000 dissociation constants $K_d$ were randomly sampled according to the uncertainty estimated in Strategy 1. For each simulated $K_d$, the corresponding apparent oligomeric state ($n_{app}$) was calculated, and the Etrue was derived. The histogram shows the resulting distribution of the 1000 simulated Etrue values. The red line indicates the mean Etrue, and dashed lines indicate the 99% confidence interval. (B) Propagation of the simulated Etrue values into Strategy 2. Each Etrue value was fixed in Eq. 5 to estimate the n app from the relationship between Eapp,se and $x_D$. The relationship used for this analysis corresponds to the AT1R dataset. The resulting $n_{app}$ values are expressed as percentages (right y-axis), while the corresponding monomer and dimer populations are shown on the left y-axis. The corresponding concentrations of monomeric and dimeric receptor species as a function of receptor surface density are shown directly below. Each point represents the mean ± 99% CI obtained from the 1000 Monte-Carlo simulations.

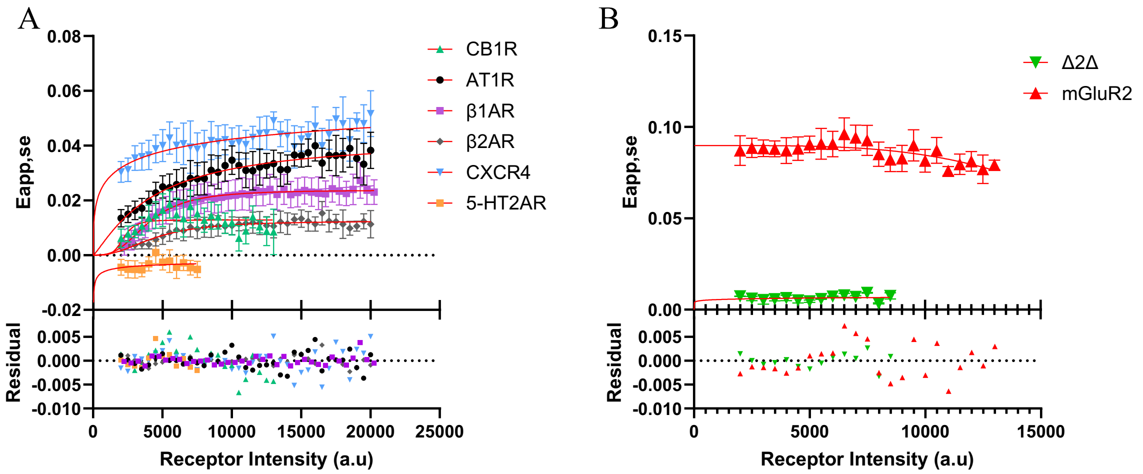


C

| Receptor | Emax | Kd (a.u) | nH |
|---|---|---|---|
| AT1R | 0.043 ± 0.008 | 4293 ± 1497 | 1.194 ± 0.487 |
| β1AR | 0.024 ± 0.002 | 4001 ± 492 | 2.875 ± 1.073 |
| β2AR | 0.013 ± 0.001 | 4409 ± 706 | 2.393 ± 0.970 |
| CXCR4 | 0.071 ± 0.100 | 3342 ± 25146 | 0.372 ± 0.717 |
| CB1R | 0.013 ± 0.001 | 2200 ± 446 | 5.219 ± 5.589 |
| 5-HT2AR | - | - | - |
| Δ2Δ | - | - | - |
| mGluR2 | 0.090 ± 0.002 | 19999 ± 6739 | 19999 |

**Figure S7: Comparative analysis of density-dependent oligomerization across class A GPCRs and controls.** (A-B) Eapp,se plotted as a function of receptor surface density (a.u.) in HEK293 cells transiently expressing SNAP-tagged class A GPCRs, including AT1R, β1AR, β2AR, CXCR4, CB1R, and 5-HT2AR (A), or mGluR2 and Δ2Δ (B). Symbols represent mean ± SEM of experimental data, and solid lines indicate nonlinear regression using the specific binding model with Hill slope (GraphPad Prism). All receptors were analyzed using a minimum of n = 3 independent experiments except Δ2Δ (n = 2). (C) Summary of apparent dissociation constants ($K_d$), Hill coefficients (nH), and maximal FRET efficiencies (Emax) derived from the fits applied to the data in A and B. Negative or nonconvergent $K_d$ values indicate the absence of detectable density-dependent population changes within the explored expression range, consistent with predominantly monomeric or dimeric receptor populations.
